# Maternal Immune Activation imprints translational dysregulation and differential MAP2 phosphorylation in descendant neural stem cells

**DOI:** 10.1101/2024.06.07.597886

**Authors:** Sandra M. Martín-Guerrero, María Martín-Estebané, Antonio J. Lara Ordóñez, Miguel Cánovas, David Martín-Oliva, Javier González-Maeso, Pedro R. Cutillas, Juan F. López-Giménez

## Abstract

Alterations induced by maternal immune activation (MIA) during gestation impact the subsequent neurodevelopment of progeny, a process that in humans, has been linked to the development of several neuropsychiatric conditions. To undertake a comprehensive examination of the molecular mechanisms governing MIA, we have devised an in vitro model based on neural stem cells (NSCs) sourced from fetuses carried by animals subjected to Poly I:C treatment. These neural progenitors demonstrate proliferative capacity and can be effectively differentiated into both neurons and glial cells. Transcriptomic, proteomic, and phosphoproteomic analyses conducted on these cellular models, in conjunction with counterparts from control treatments, revealed discernible shifts in the expression levels of a specific subset of proteins implicated in neuronal function. Noteworthy, we found an absence of congruence between these alterations at the transcriptomic level, suggesting that differences in protein translation contribute to the observed dysregulation. Furthermore, the phosphoproteomic data highlighted a discernible discrepancy in the basal phosphorylation of proteins between differentiated cells from both experimental groups, particularly within proteins associated with cytoskeletal architecture and synaptic functionality, notably those belonging to the MAP family. Observed alterations in MAP phosphorylation were found to potentially have functional consequences as they correlate with changes in neuronal plasticity and the establishment of neuronal synapses. Our data agrees with previous published observations and further underscore the importance of MAP2 phosphorylation state on its function and the impact that this protein has in neuronal structure and function.

## INTRODUCTION

It is widely accepted that environmental stressors during pregnancy may affect the normal development of the central nervous system, leading to the manifestation of different neuropsychiatric diseases such as schizophrenia, autism, or bipolar disorders. Among the environmental factors that can influence fetal neurodevelopment from the post-conception period are the infectious processes experienced by pregnant females during gestation [1]. Intriguingly, some behavioral manifestations corresponding to the pathological phenotype appear during late adolescence or early adulthood in the offspring, as observed in the case of schizophrenia [2, 3].

Therefore, several preclinical models based on experimental animals have been developed in order to explore the biological mechanisms resulting from prenatal infections that act as primers for these neurodevelopmental diseases [4, 5]. Originally these models consisted essentially in exposing animals, preferentially rodents, to the infectious pathogen, namely bacteria or virus, during the early stages of pregnancy. In the case of viruses, alternative methodologies exist where the pathogen is substituted by chemical activating agents that induce an equivalent pro-inflammatory immune response in the mother. One such example is polyinosinic : polycytidylic acid or Poly (I:C), a synthetic analog of double-stranded RNA. When administered to animals, it can mimic the acute phase of viral infection, resulting in a comprehensive range of behavioral and cognitive abnormalities that become apparent only when the offspring reach late adolescence or early adulthood.

Previous studies conducted with Poly (I:C) in pregnant mice have demonstrated that the determinant factor responsible for neurodevelopmental disturbances after an infectious process is the acute generation of pro-inflammatory cytokines, including various interleukins and TNF-α, rather than the presence of the pathogen itself [6]. Therefore, this maternal immune activation or MIA during pregnancy is sufficient for inducing a range of disease phenotypes in offspring. More precisely, it was demonstrated that the participation of IL-6 in the maternal immune response is necessary since the elimination of this cytokine through genetic engineering techniques deprived animals of the behavioral deficits associated with MIA in the adulthood [7]. Similarly, the necessary participation of IL-17A in the neurodevelopment process following Poly I:C treatment was also demonstrated later on [8, 9]. Further investigations also determined the gestational time window during which maternal immune activation is most effective in inducing subsequent behavioral alterations in adult descendants. This aligns to the early period of pregnancy, specifically gestational day 9 in mice, which corresponds to the first to second trimester in humans [6].

Overall, these preclinical studies supported the neurodevelopmental hypothesis of schizophrenia and other mental diseases associated with infectious processes in the maternal host during early stages of pregnancy. This is evidenced by the pathological behavioral phenotype observed in offspring, which notably is not accompanied by significant anatomical alterations in their brains [2, 3, 5]. Consequently, this leads to conjecture that all these neurodevelopmental disturbances initiated by MIA, which are responsible for the pathological phenotypes observed in descendants, may be regarded as transcriptomics and epigenomics alterations occurring during the fetal neurodevelopment. In this sense, a systematic review recently published compiles data from multiple investigations assessing quantitative changes in gene and protein expression as well as epigenetic markers obtained in brain samples from descendants of MIA models [10]; the evaluation of a total of 118 studies concluded that changes in candidate genes were mainly associated with three key functional areas, including immune and stress responses, neurotransmission and neuronal signaling, and neurodevelopment. However, the precise molecular changes and mechanisms of this process remain elusive.

Here, to delve deeper into the biological mechanisms implicated in the MIA effects on neurodevelopment processes and investigate the alterations that persist in cultured neural lineages derived from their differentiation, we have generated neural stem cell lines from fetuses carried by pregnant mice treated with Poly I:C. Once stablished, we characterized these models using transcriptomics, proteomics and phosphoproteomics approaches. Our findings suggest that Poly I:C meditated immune activation during early gestation induces modest translational dysregulation, but interestingly, this treatment result in comparatively large alterations in the phosphorylation of proteins involved in processes critical for the organization of neuronal structure. Of relevance, we find these such changes have potential functional consequences in increasing neuronal spine density and while decreasing synaptogenesis.

## MATERIALS AND METHODS

### Drugs

Poly (I:C) (Polyinosinic-polycytidylic acid sodium salt, lots #059M4167V and #0000110446) were purchased from Sigma-Merck.

### Animals

C57BL/6J mice were purchased from Charles River (Spain). All the experimental work carried out in this study followed the ethical guidelines from Directive 2020/569/EU of the European Parliament on the protection of animals used for scientific purposes. Animals were housed in a controlled-temperature/humidity environment (22 ± 1 °C, 60-70 % relative humidity) in individual cages with a 12h light/dark cycle and food and water available at libitum. According to the protocol established by the experimental animal unit of the IPBLN-CSIC, mice were allowed to acclimatize to housing room for a minimum of 5 days after their reception in the animal facility.

### Maternal Immune Activation (MIA) experiments

After mouse mating, female pregnancies were determined by visualizing seminal plug (E0.5) and by body weight increase. Poly(I:C) treatment was administered intraperitoneally on day E9.5. Various doses of Poly(I:C) were tested, starting at 20 mg/kg and gradually decreasing to 5 mg/kg due to the elevated animal mortality observed at the highest doses. The experimental control group was treated in parallel by injecting mice with an equivalent volume of saline. Blood samples from treated animals were taken 2.5h after treatment by mandibular sinus venipuncture in order to test later IL-6 levels in serum. Pregnant females were submitted to cesarean surgery on day E12.5 to facilitate fetal dissection, collecting the pooled fetal tissue as the initial biological material for obtaining neural stem cells (NSC). Three different NSC lines were obtained from each experimental condition, i.e., Poly I:C and Saline treatments, coming from independent pregnant mice. Blood samples from females were also taken by cardiac puncture to verify IL-6 levels in serum at this time point. The temperature, weight, and any signs of sickness in pregnant females were monitored throughout the entire process. Additionally, recommendations on experimental practices in MIA experiments, published elsewhere, were followed [11, 12]. IL-6 serum levels were measured using an ELISA kit (88-7064, Invitrogen). The assay was performed following the manufactureŕs instructions and guidelines. Briefly, Nunc MaxiSorpTM flat-bottom 96-well-plates (Thermo Scientific) were coated with anti-mouse IL-6 capture antibody overnight at 4°C. Plates were washed with PBS + 0.05% Tween20 and incubated with ELISA/ELISPOT Diluent for 1h to block non-specific binding sites. Plasma samples diluted in ELISA/ELISPOT Diluent were incubated overnight at 4°C. A standard curve was generated using the IL-6 standard provided. After washing, plates were incubated with biotin-conjugated anti-mouse IL-6 antibody for 1h, then washed and incubated with Avidin-HRP for 30min. Plates were washed and finally incubated with Tetramethylbenzidine (TMB) Substrate Solution for 15min. The reaction was stopped with 2N H_2_SO_4_ and the signal was measured on a plate reader at 450 nm.

### Cell culture

Adherent neural stem cell (NSC) lines from the mouse forebrain were obtained following the methodology previously described [13]. Briefly, the fetus dorsal forebrain at E12.5 was dissected, and the obtained tissue was disaggregated in order to get a cell suspension. These cells were then transferred to cell-culture plastic plates that had been previously treated with poly-L-lysine 10 μg/ml (Sigma-Merck) to enhance cellular adhesion and promote monolayer cultures. NSCs were maintained and expanded in NS expansion medium composed as follows: DMEM/Ham’s F-12 media with l-glutamine (Gibco), Glucose solution 29mM (Signa-Merck), MEM nonessential amino acids 1x (Gibco), Penicillin/Streptomycin 1x (Gibco), HEPES buffer solution 4.5 mM (Cytiva), BSA solution 0.012% (Gibco), 2-mercaptoethanol 0.05 mM (Gibco), N2 supplement 1x (Gibco), B27 supplement 1x (Gibco), murine epidermal growth factor (EGF) 10 ng/ml (PeproTech) and human fibroblast growth factor (FGF-2) 10 ng/ml (PeproTech). To induce differentiation of NSCs into various neural cell lineages, the expansion medium was replaced with the following differentiation medium: Neurobasal plus DMEM/Ham’s F-12 media with l-glutamine 1:1 (Gibco), N2 supplement 1x (Gibco), B27 supplement 1x (Gibco), Glutamax 1x (Gibco) and 2-mercaptoethanol 0.05 mM (Gibco). Cells were grown and maintained at 37 °C-5% CO2 under a humidified atmosphere.

### Immunocytochemistry

Cells were seeded on glass-coverslips pretreated with poly-L-lysine 50 μg/ml plus laminin 3.3 μg/ml (Sigma-Merck) and fixed with 4% paraformaldehyde before immunolabeling. Primary antibodies used were anti-Nestin 1:100 (Abcam, ab254048), anti-Sox2 1:100 (Abcam, ab97959), anti-MAP2 1:200 (SIGMA, M3696), anti-β-III tubulin 1:100 (R&D Systems, MAB1195-SP), anti-GFAP 1:100 (Cell Signaling, #3670) and anti-VGLUT2 1:200 (Abcam, ab79157). Secondary antibodies were anti-mouse IgGAlexaFluor 488 1:1000 (Abcam, ab150105) and anti-rabbit IgGAlexaFluor 594 1:1000 (Abcam, ab150076). Fluorescent cell nuclei staining was performed with Hoechst 33342 (Invitrogen) according to the manufacturer instructions. Fluorescent images of cells were acquired using an Olympus Ix81 epifluorescence inverted microscope or a Leica TCS SP5 confocal microscope. Image processing was conducted with Image J software.

### Neuritogenesis and spinogenesis analysis

NSCs were seeded into 6-well plates pre-treated with poly-L-lysine 10 μg/ml plus laminin 3.3 μg/ml. After 7 days of differentiation, the complexity of dendritic arborization was assessed in images acquired through phase-contrast microscopy using a 40x objective, considering a population of 10-15 neurons. The Sholl analysis tool, included in ImageJ software, was utilized to estimate dendritic complexity. This analysis involves establishing concentric circles from the center of the neuronal body and counting the number of intersections these circles make with the various dendritic processes emerging from the neuron soma. The resulting data can be graphed, depicting the number of intersections in relation to the distance from the center of the neuron. The maximum number of intersections per neuron obtained from this graph was considered to compare and average the results from different experiments. Spinogenesis were evaluated quantifying the dendritic spine density in images acquired through phase-contrast microscopy using a 63x objective and Leica DMi8 microscope. The number of dendritic spines were quantified in 20 μm segments of dendrites using ImageJ software.

### Synaptogenesis

The quantitative evaluation of established synapses among in vitro cultured neurons was conducted using a rabies virus monosynaptic tracing methodology, which involved the utilization of genetically modified rabies viruses pseudotyped to infect a specific set of neurons expressing the appropriate virus receptor [14]. The engineered version of the rabies virus harbors in its genome a gene encoding a fluorescent protein, which replaces the original gene corresponding to the capsid glycoprotein. This modification enables rabies virus particles to be transmitted to presynaptic neurons through neurite terminals in a retrograde manner. The absence of the native capsid glycoprotein gene prevents the further transmission of the virus to subsequent neurons, limiting the infection to a unique presynaptic level. The generation of genetically engineered rabies virus was performed following the protocol previously described [15]. Plasmids pSADdeltaG-F3 (32634), pcDNA-SADB19N (32630), pcDNA-SADB19P (32631), pcDNA-SADB19L (32632), pcDNA-SADB19G (32633) and pBOB-synP-HTB (30195) were acquired from Addgene. The original pSADdeltaG-F3 plasmid was modified by subcloning the encoding sequence of mCherry protein between NheI and SacII restriction sites. HEK 293T-TVA 800, B7GG and BHK-EnvA cell lines were generously provided by Edward Callaway from The Salk Institute for Biological Studies (California, USA). Recombinant lentivirus production was conducted according to standard protocols from Addgene using pBOB-synP-HTB in combination with psPAX2 and pCAG-VSVG as packaging and enveloping helper plasmids respectively. To visualize monosynaptic tracing with rabies virus in cultured neurons, NSCs were seeded in 6-well tissue culture plates pre-treated with poly-L-lysine (10 µg/ml) plus laminin (3.3 µg/ml) at a density of 200,000 cells per well. Once the differentiation process began (day 0), on day 3 cells were infected with the lentivirus to induce neurons to express the TVA receptor along with GFP and rabies capsid glycoprotein. Subsequently, on day 5, cells were infected with the recombinant rabies virus, and finally, on day 7, image acquisition from living cells was performed using a Leica DMI8 epifluorescence microscope with a 10x objective after staining cell nuclei with Hoechst 33342. Cells were illuminated to visualize fluorescent signals corresponding to GFP, mCherry, and Hoechst 33342 emission wavelengths, namely green, red, and blue, respectively. Thus, the blue channel showed the total population of cells within the microscopy field, whereas the green channel corresponded to those cells expressing the TVA receptor and therefore susceptible to infection by the recombinant rabies virus. Finally, the red channel showed those cells infected by the rabies virus. Within this population of cells expressing the mCherry protein, there were cells displaying both GFP and mCherry fluorescent signals, as well as cells emitting only the mCherry signal. These latter cells represent neurons infected by the rabies virus through monosynaptic connections (Supplemental Figure 4 showing the 3 microscopy fields and pointing out neurons not co-expressing green and red signals). Image analysis to quantify the different cell populations was performed using FIJI software, employing a macro specifically created for this purpose. This macro consisted of Gaussian blur sigma filtering applied prior to image segmentation, which was based on autothresholding of cell nuclei stained with Hoechst 33324 reagent. Several morphological parameters could be adjusted to optimize the segmentation process, such as minimum and maximum cell size, minimum circularity, and thresholds of GFP and mCherry minimum and maximum fluorescence signals. To evaluate the extent of synapses formed among neurons present in the analyzed microscopy field, we considered the ratio between the total number of cells displaying only the mCherry signal and the total number of cells displaying both GFP and mCherry signals simultaneously.

### Immunoblotting

Samples were heated at 65°C for 15 min and subjected to SDS-polyacrylamide gel electrophoresis analysis using 4 to 12% bis-Tris gels (NuPAGE, Invitrogen) and MOPS buffer. After electrophoresis, proteins were transferred onto nitrocellulose membranes that were incubated in a solution of 5% non-fat milk and 0.1% Tween-20 in Tris-buffered saline at room temperature on a rotating shaker for 2 hours to block nonspecific binding sites. Afterwards, membranes were incubated overnight with a rabbit anti-MAP2 polyclonal antibody 1:1000 (SIGMA M3696). Protein antibody interactions were detected using horseradish peroxidise-linked anti-rabbit IgG 1:10000 (Abcam ab6721). Immunoblots were developed by application of enhanced chemiluminescence solution (ECL, Cytiva). Immunoblots were stripped subsequently and re-blotted with anti-actin antibody 1:5000 (Hypermol, clone262) to assess the amount of loaded protein.

### Sample preparation for proteomics and phosphoproteomics analysis

Mass spectrometry experiments were performed as described previously [16] with some modifications. Cultured cells on 100 mm dishes were washed with ice-cold PBS supplemented with phosphatase inhibitors (1mM Na3VO4 and 1mM NaF). Cells were scrapped and lysed with a Urea buffer (8M urea in 20mM in HEPES, pH 8.0 supplemented with 1mM Na3VO4, 1mM NaF, 1mM Na2H2P2O7 and 1mM sodium β-glycerophosphate) and lysates were then sonicated using a sonicator Vibra-Cells (Sonics Materials) at 20% of intensity, 3 cycles of 10 seconds. Samples were then centrifuged at 13,000 rpm for 10 min at 5°C. Supernatant was transferred in 1.5mL Protein Lo-bind tube (Eppendorf). Protein concentration was determined using BCA Protein Assay Kit. For phosphoproteomics analysis, 110 µg of extracted proteins in a volume of 200 ul were reduced with dithiothreitol (DTT, 10 mM) for 1 h at 25 °C, and alkylated with Iodoacetamide (IAM, 16.6 mM) for 30 min at 25 °C. Trypsin beads were equilibrated by three washes with 20 mM HEPES; pH 8.0. Then, samples were diluted with 20 mM HEPES (pH 8.0) to a final concentration of 2 M urea and digested with equilibrated trypsin beads (50% slurry of TLCK-trypsin) overnight at 37 °C. After that, trypsin beads were removed by centrifugation (2000×g for 5 min at 5 °C) and samples were transferred to 96 well plates and acidified by adding TFA to a final concentration of 0.1%. Peptide solutions were desalted and subjected to phosphoenrichment using the AssayMAP Bravo (Agilent Technologies) platform. For desalting, protocol peptide clean-up v3.0 was used. Reverse phase S cartridges (Agilent, 5 μL bed volume) were primed with 250 μL 99.9% acetonitrile (ACN) with 0.1%TFA and equilibrated with 250 μL of 0.1% TFA at a flow rate of 10 μL/min. The samples were loaded (770 μL) at 20 μL/min, followed by an internal cartridge wash with 250 μL of 0.1% TFA at a flow rate of 10 μL/min. Peptides were then eluted with 105 μL of 1M glycolic acid with 50% ACN, 5% TFA and this is the same buffer for subsequent phosphopeptide enrichment. Following the Phospho Enrichment v 2.1 protocol, phosphopeptides were enriched using 5µl Assay MAP TiO2 cartridges on the Assay MAP Bravo platform. The cartridges were primed with 100µl of 5% ammonia solution with 15% ACN at a flow rate of 300 μL/min and equilibrated with 50 μL loading buffer (1M glycolic acid with 80% ACN, 5% TFA) at 10 μL/min. Samples eluted from the desalting were loaded onto the cartridge at 3 μL/min. The cartridges were washed with 50 μL loading buffer and the phosphorylated peptides were eluted with 25 μL 5% ammonia solution with 15% ACN directly into 25 μL 10% formic acid. Phosphopeptides were lyophilized in a vacuum concentrator and stored at -80°C. For proteomics analysis, 30 μg of protein were digested and acidified as described above. Peptide solutions were desalted using the AssayMAP Bravo (Agilent Technologies) platform. Cartridges were primed, equilibrated, loaded and washed as described above and peptides were eluted with 105 μL of 70/30 ACN/ H2O + 0.1% TFA. Eluted peptide solutions were dried in a SpeedVac vacuum concentrator and peptide pellets were stored at −80 °C.

### LC-MS/MS Analysis

Peptides were re-suspended in 20 µL of reconstitution (97% H2O, 3% ACN, 0.1% TFA, 50fmol/µl-1 enolase peptide digest) and sonicated for 5 minutes at RT. Following a brief centrifugation, 2μl was loaded onto a LC-MS/MS system. This consisted of a nano flow ultra-high pressure liquid chromatography system UltiMate 3000 RSLC nano (Dionex) coupled to a Q Exactive Plus using an EASY-Spray system. The LC system used mobile phases A (3% ACN: 0.1% FA) and B (100% ACN; 0.1% FA). Peptides were loaded onto a μ-pre-column and separated in an analytical column. The gradient: 1% B for 5 min, 1% B to 35% B over 90min, following elution the column was washed with 85% B for 7 min, and equilibrated with 3% B for 7min, flow rate of 0.25 µL/min. Peptides were nebulized into the online connected Q-Exactive Plus system operating with a 2.1s duty cycle. Acquisition of full scan survey spectra (m/z 375-1,500) with a 70,000 FWHM resolution was followed by data-dependent acquisition in which the 15 most intense ions were selected for HCD (higher energy collisional dissociation) and MS/MS scanning (200-2,000 m/z) with a resolution of 17,500 FWHM. A 30s dynamic exclusion period was enabled with an exclusion list with 10ppm mass window. Overall duty cycle generated chromatographic peaks of approximately 30s at the base, which allowed the construction of extracted ion chromatograms (XICs) with at least ten data points.

### Peptide and protein identification and quantification

Peptide identification from MS data was automated using a Mascot Daemon (v2.8.0.1) workflow in which Mascot Distiller generated peak list files (MGF) from RAW data, and the Mascot search engine matched the MS/MS data stored in the MGF files to peptides using the SwissProt Database restricted to Mus musculus (SwissProt_2021_02.fasta, 17080 sequences). Searches had an FDR of ∼1% and allowed 2 trypsin missed cleavages, mass tolerance of ±10 ppm for the MS scans and ±25 mmu for the MS/MS scans, carbamidomethyl Cys as a fixed modification and oxidation of Met, PyroGlu on N-terminal Gln and phosphorylation on Ser, Thr, and Tyr as variable modifications (phosphorylation was only considered for searches performed in phosphoproteomics data). Pescal was used for label free quantification of the identified peptides [17]. The software constructed XICs for all the peptides identified in at least one of the LC-MS/MS runs across all samples. XIC mass and retention time windows were ±7 ppm and ±2 min, respectively. Quantification of peptides was achieved by measuring the area under the peak of the XICs. Individual peptide intensity values in each sample were normalized to the sum of the intensity values of all the peptides quantified in that sample. Phosphoproteomics and proteomics data was processed and analysed using a bioinformatic pipeline developed in a R environment (https://github.com/CutillasLab/protools2/). The normalised data was centered, log2 scaled and 0 values were inputted using the minimum feature value in the sample minus one. Then, p-values to assess statistical differences between comparisons were calculated using LIMMA [18], and then adjusted for FDR using the Benjamini-Hochberg procedure. Differences were considered statistically significant when p-values <0.05 and FDR<0.25.

### Sequencing and data processing of transcriptomic data

NSCs were plated in 100 mm culture dishes and washed with ice-cold PBS. Cellular pellets from either non-differentiated NSCs or cells differentiated for 7 days were collected and processed for transcriptomics approaches. Both, library preparation and Illumina sequencing were carried out at the IPBLN Genomics Facility (CSIC, Granada, Spain). Total RNA integrity was verified by Bioanalyzer RNA 6000 Nano chip electrophoresis (Agilent Technologies). Every RNA sample showed a RIN value above 8.5. RNA-seq libraries were prepared using Illumina stranded mRNA Prep Ligation kit (Illumina®) from 200 ng of input total RNA. Quality and size distribution of PCR-enriched libraries were validated through Bioanalyzer High Sensitivity DNA assay and concentration was measured on the Qubit® fluorometer (Thermo). Final libraries were pooled, in an equimolecular manner, and then diluted and denatured as recommended by Illumina NextSeq® 500 library preparation guide. The 72×2 nt paired-end sequencing was conducted on a NextSeq® 500 sequencer with a final output of 48 Gbp and a quality score (Q30) of 88.5 %. miARma-Seq pipeline was used to analyse transcriptomic samples [19, 20]. Firstly, raw data were evaluated using FastQC software to analyze the quality of the reads (http://www.bioinformatics.babraham.ac.uk/projects/fastqc). Subsequently, after sample filtering, we obtained a mean of 49% in GC content and an average of 23,636,605 reads per sample. No adapter accumulation and bad quality reads (q<30) were found. Afterwards, the number of reads per sample were homogenized using Seqtk software (https://github.com/lh3/seqtk). In the second step, miARma-Seq aligns all sequences using STAR [21],resulting in a 93.28% of properly aligned reads. With this aim, we use the Mus musculus Gencode version M31 genome-build: mmGRCm39 was used. After that, featureCounts software [22] was used to assign sequence reads to genes by using reference gene annotation that was obtained from Gencode from the same assembly and genome build. Expression analysis was carried out by edgeR package [23]. Low expressed genes were removed, and remaining genes were normalized by the trimmed mean of M-values (TMM) method [24]. The normalized gene expression (TMM) was log2 scaled and then, statistical differences between comparisons were calculated using LIMMA, and then adjusted for FDR using the Benjamini-Hochberg procedure. Differences were considered statistically significant when p-values <0.05 and FDR<0.25.

### Statistics

Statistical analysis was carried out in Excel or in R (v4.3.1) using base functions or the ggpubr package (https://CRAN.R-project.org/package=ggpubr). When not specified, statistical differences were evaluated using unpaired T-Student test. Gene Ontology analysis of genes/proteins differentially expressed or phosphorylated was performed using the clusterProfiler package (https://bioconductor.org/packages/release/bioc/html/clusterProfiler.html). Data was visualized using the ggplot2 package (https://cran.r-project.org/web/packages/ggplot2/index.html).

## RESULTS

### Generation of neural stem cell lines from fetuses carried by mice exposed to MIA treatment

Neural stem cells (NSCs) were obtained from fetuses at E12.5 to establish adherent proliferative cell lines. Pregnant females were treated either with Poly(I:C) or saline three days before, corresponding to E9.5. This timing aligns with gestation day 9, a critical point when MIA treatment has been shown to induce significant behavioral alterations in the offspring [6, 11]. Of note, due to the high mortality rate observed when treating animals with the Poly(I:C) dose described previously (i.e., 20 mg/kg i.p. [7, 12, 25]), we had to gradually reduce the dose and test different Poly(I:C) batches until reaching 5 mg/kg i.p. Because late behavioral analysis of offspring was not feasible in this experimental protocol as a phenotypic indicator of MIA during pregnancy, we assessed maternal levels of IL-6 in serum 2.5 hours post-injection and significant fluctuations in body temperature as reliable indicators of MIA leading to disease-like phenotypes in the descendants [12]. Supplementary Figure 1A summarizes the protocol indicating the time points when body temperature and blood samples were taken, from the moment of the Poly(I:C) injection to the cesarean surgery. Substantial changes were observed in three of the four surviving animals treated with Poly(I:C) affecting both phenotypical parameters. Specifically, there was a considerable hypothermia 2.5 h post-injection (Supplementary Figure 1B) accompanied by a robust increase of IL-6 concentration in serum to some 10,000 pg/ml (Supplementary Figure 1C). Both body temperature and IL-6 levels in serum returned to levels similar to the saline group three days after Poly(I:C) administration.

Fetal forebrain tissue dissected from each of these pregnant females was pooled to generate three independent neural stem cell lines. A similar procedure was followed with animals treated with saline, serving as the experimental control group. Adherent cell lines obtained from both experimental groups proliferated after subsequent passages and expressed markers associated with pluripotency such as Sox-2 and Nestin (Supplementary Figure 2). Next, we determined the cell culture conditions to prompt the differentiation of NSCs into distinct neural lineages, specifically neurons and glia. This was achieved by replacing the expansion growth medium with differentiation medium. Under these experimental conditions, NSCs underwent phenotypic transformation over time, resulting in cells that could be morphologically identified as neurons and astrocytes (Supplementary Figure 3). The differentiation process was distinctly observable through phase-contrast microscopy after 4-5 days. Additional immunocytochemistry experiments were conducted to identify neuronal and glial cells using specific markers such as antibodies recognizing β-III tubulin and GFAP respectively (Figure 1). Furthermore, in the case of neurons, they showed a preferential immunopositivity for VGLUT2, thereby indicating their glutamatergic neurochemical nature (Figure 1). No differences were observed between the two experimental groups, i.e., NSCs obtained from Poly I:C treatments versus Saline, in terms of differentiation into neurons and glial cells, as well as regarding labeling with immunocytochemistry markers. We thus stablished a robust protocol for the generation of NSCs.

**Figure 1.**
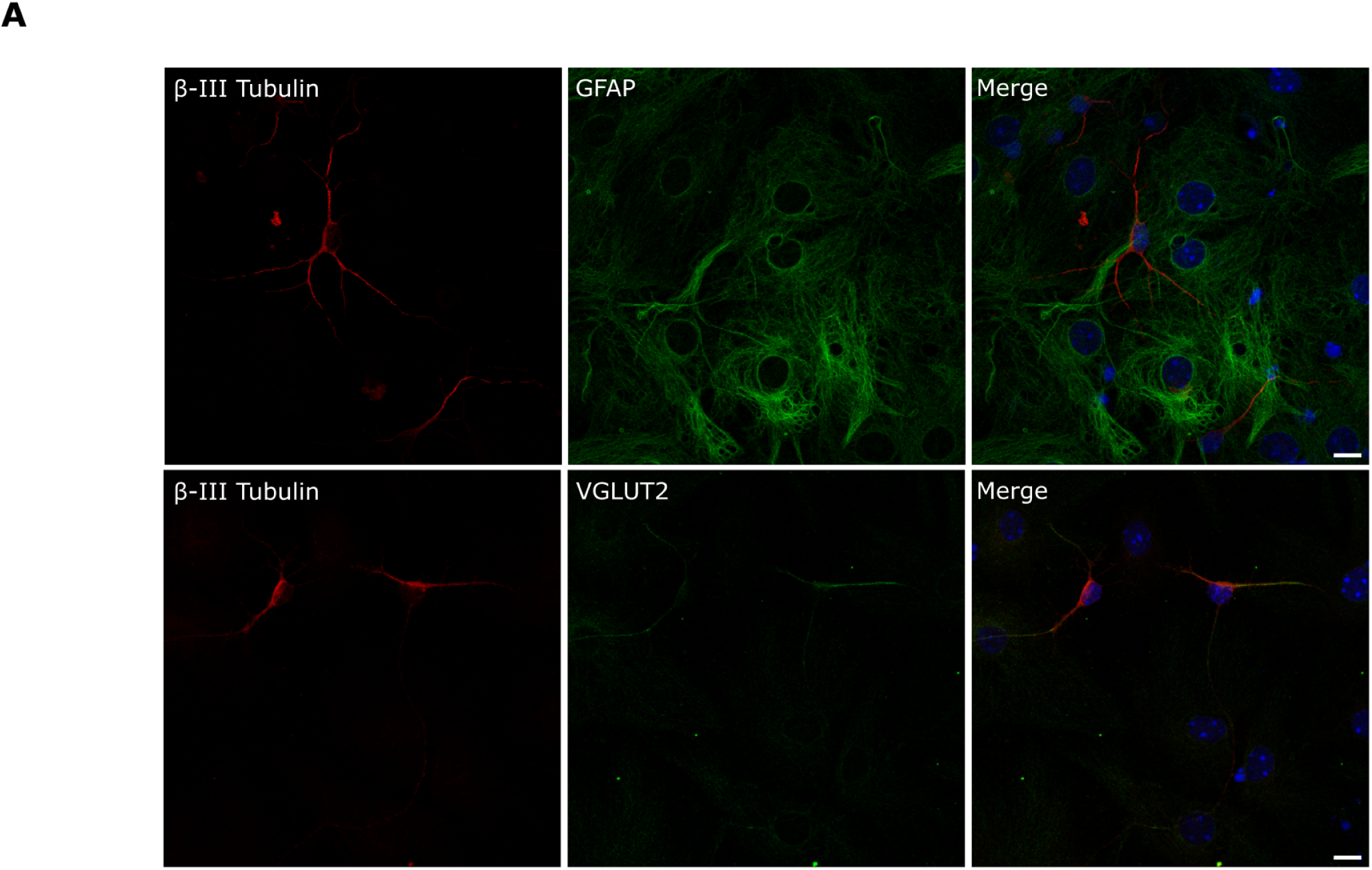
Representative images of NSCs cell lines after 5 days of differentiation into distinct neural lineages, specifically neurons and glia. Immunocytochemistry experiments were conducted to identify neuronal and glial specific markers such as ß-III Tubulin and GFAP respectively. VGLUT2 staining was performed to detect glutamatergic neurons. Scale bar: 10 µm.

### Transcriptomics and proteomics analysis from Poly I:C and Saline NSCs

To characterise our NSCs models molecularly, cells from Poly I:C and Saline groups were cultured and differentiated for 7 days and then subjected to transcriptomics and proteomics analysis. Non-differentiated NSCs were also subjected to the same analysis in parallel. The analysis of transcriptomic data showed that the differentiation of Saline NSCs modified the expression of 4409 genes (2990 increased and 1419 decreased, Figure 2A left volcano plot). The differentiation of Poly I:C NSCs also significantly modified the expression of 5877 genes (3530 increased and 2347 decreased, Figure 2A right volcano plot). The analysis of protein expression from proteomics data showed that differentiation of Saline NSCs significantly increased the expression of 389 proteins and decreased the expression of 569 proteins (Figure 2B); whereas the differentiation of Poly I:C NSCs increased and decreased the expression of 336 and 649 proteins, respectively (Figure 2B). We next performed Gene Ontology analysis for the significant genes and proteins modified in each group. These data showed that the differentiation of NSCs into neurons and glial cells significantly reduced the expression of genes associated with cell division and translation processes, and it increased the expression of genes related with cilium processes and axoneme assembly (Figure 2C), as expected. All the significant Gene Ontology terms identified in both groups are indicated in Supplementary Table 1A-B. As observed for transcriptomic analysis, Gene Ontology analysis from proteomics data also showed that the differentiation process of both types of NSCs decreased the expression of proteins related with RNA processes (Figure 2D, Supplementary Table 1C-D). In addition, we also observed an increase in the expression of proteins related with cytoskeleton organization and actin-based processes (Figure 2D). To further evaluate the differentiation process of NSCs in both groups, we analysed common markers of stem cells and neuronal markers for differentiated neurons and glia. As stem cell markers we evaluated the expression levels of protein Ki-67 (Mki67), DNA replication licensing factor MCM2 (Mcm2), the RNA-binding protein Musashi-2 (Msi2), Nestin (Nes) and the transcription factor Sox2. For neuronal and glial markers, we evaluated the expression of the Neuronal migration protein doublecortin (Dcx), Cadherin-2 (Cdh2), the Kinesin light chain 1 (Klc1), Protein S100-B (S100b), the SNARE-associated protein SNAPIN (Snapin), the Excitatory amino acid transporter 1 (Slc1a3), Synapse-associated protein 1 (Syap1) and Synaptopodin-2 (Synpo2) (Figure 2E-F). As expected, the differentiation of NSCs decreased the expression of stem cell markers and increased the expression of neuronal/glial markers in transcriptomics and proteomics analysis (Figure 2E-F). After confirming that differentiation of NSCs induced changes in both type of cell lines, we evaluated the presence of differences between Saline and Poly I:C before and after the differentiation process. Interestingly, the analysis of transcriptomics data showed that no significant changes were present when analysing Poly I:C NSCs compared to Saline NSCs (Figure 3A). In the case of differentiated NSCs, only one gene was significantly reduced in Poly I:C compared to Saline (Figure 3A), Gasdermin-D (Gsdmd), involved in the response to pathogens [26]. In agreement with transcriptomics analysis, the analysis of protein expression levels also revealed no significant differences between NSCs from Poly I:C and Saline groups (Figure 3B). Unexpectedly, in contrast to transcriptomics data, differentiated Poly I:C NSCs presented altered expression levels of 69 proteins compared to differentiated Saline NSCs (Figure 3B). We found 47 proteins significantly downregulated and 22 significantly upregulated in differentiated Poly I:C NSCs compared to differentiated Saline NSCs (Figure 3B-C). In addition, we observed a distinct pattern of protein expression between the Saline and Poly I:C groups; specifically, while protein expression decreased or increased as a consequence of the differentiation of Saline NSCs, this trend was not observed in Poly I:C NSCs (Figure 3C-D). For example, this disparity can be observed in the case of Anapc1 expression, i. e., it decreased upon differentiation of Saline NSCs, whereas this effect was not observed in Poly I:C NSCs. In other cases, this decrease was higher after the differentiation of Poly I:C NSCs in comparison to differentiated Saline NSCs (e.g. Gatad2b, Figure 3C-D). Also, an opposite pattern could be observed when analysing proteins which levels were increased after the process of differentiation in Saline NSCs (Figure 3C), that is, the differentiation of Poly I:C NSCs did not show significant increase in their expression. An illustration of this circumstance is Histone deacetylase 1 (Hdac1), which is involved in neuronal differentiation [27]; it was only significantly increased after the differentiation of Saline NSCs (Figure 3C-D). Similarly, we observed this trend in the protein levels of the neuronal SNARE-binding protein Snapin, important for endocytic trafficking in neurons [28]; the differentiation of Poly I:C NSCs did not increase Snapin expression as occurred for the Saline group (Figure 3D). A comparable pattern can be observed in other proteins as observed in Figure 3C and D.

**Figure 2.**
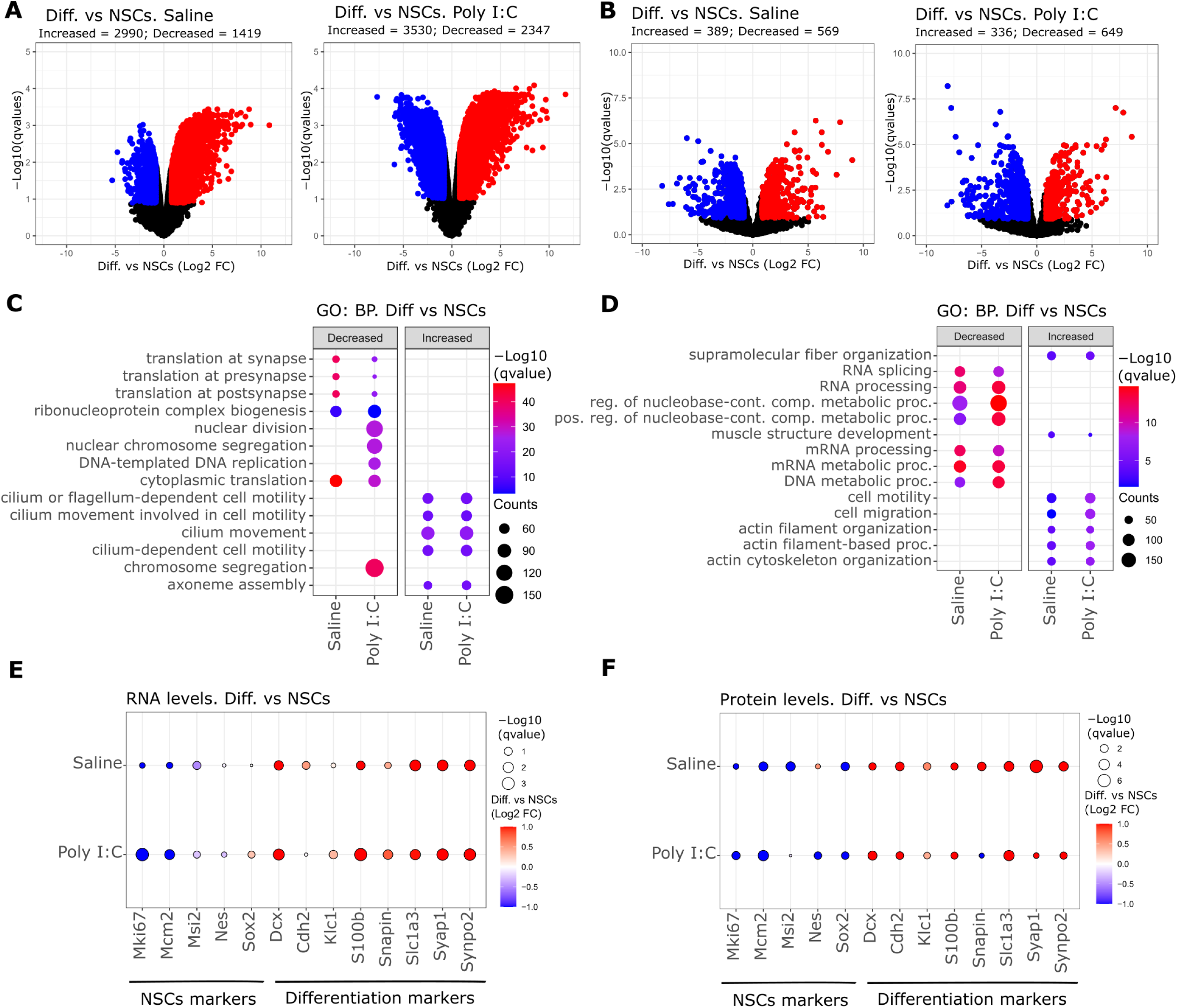
Transcriptomics and proteomics analysis of Saline and Poly I:C NSCs after differentiation process. **A, B.** Volcano plots showing transcriptomics (**A**) and proteomics (**B**) data from Saline and Poly I:C after differentiation of NSCs. X-axis shows relative change expressed as Log2 Fold Change (FC) of differentiated cells versus NSCs in each cell type. Y-axis show statistical significance expressed as -Log10 qvalues (FDR). Red and blue dots correspond to genes or proteins that changed significantly respect to NSCs cells (qvalues<0.25 & pval <0.05 and Log2 FC<-0.8 or Log2 FC> 0.8, respectively); black dots represent genes or proteins that did not match with the filtering criteria (FDR>0.25 & pval > 0.05, Log2 FC > −0.8 or Log2 FC <0.8). **C, D.** Gene Ontology (GO) analysis for Biological Processes performed on the significant genes (**C**) and proteins (**D**) from the volcano plots in A and B. Plots show the top 5 most significant terms in each comparison and cell line. Decreased and increased indicates GO terms which genes or protein were found with significant decreased or increased levels (blue and red dots in volcano plots). The size of the dots represents the number of genes or proteins identified for each term, while the gradient colour scale represents the level of significance, indicated as -Log10 qvalue. **E, F.** Dot plots showing the relative levels of genes (RNA levels, **E**) and proteins (**F**) of neural stem cell markers (NSCs) and differentiation markers in Saline and Poly I:C differentiated cells compared to NSCs. The size of the dots indicated the level of significance for each comparison, indicated as -Log10 qvalues (FDR). The colour of the dots indicates the relative change expressed as Log2 FC of differentiated cells versus NSCs.

**Figure 3.**
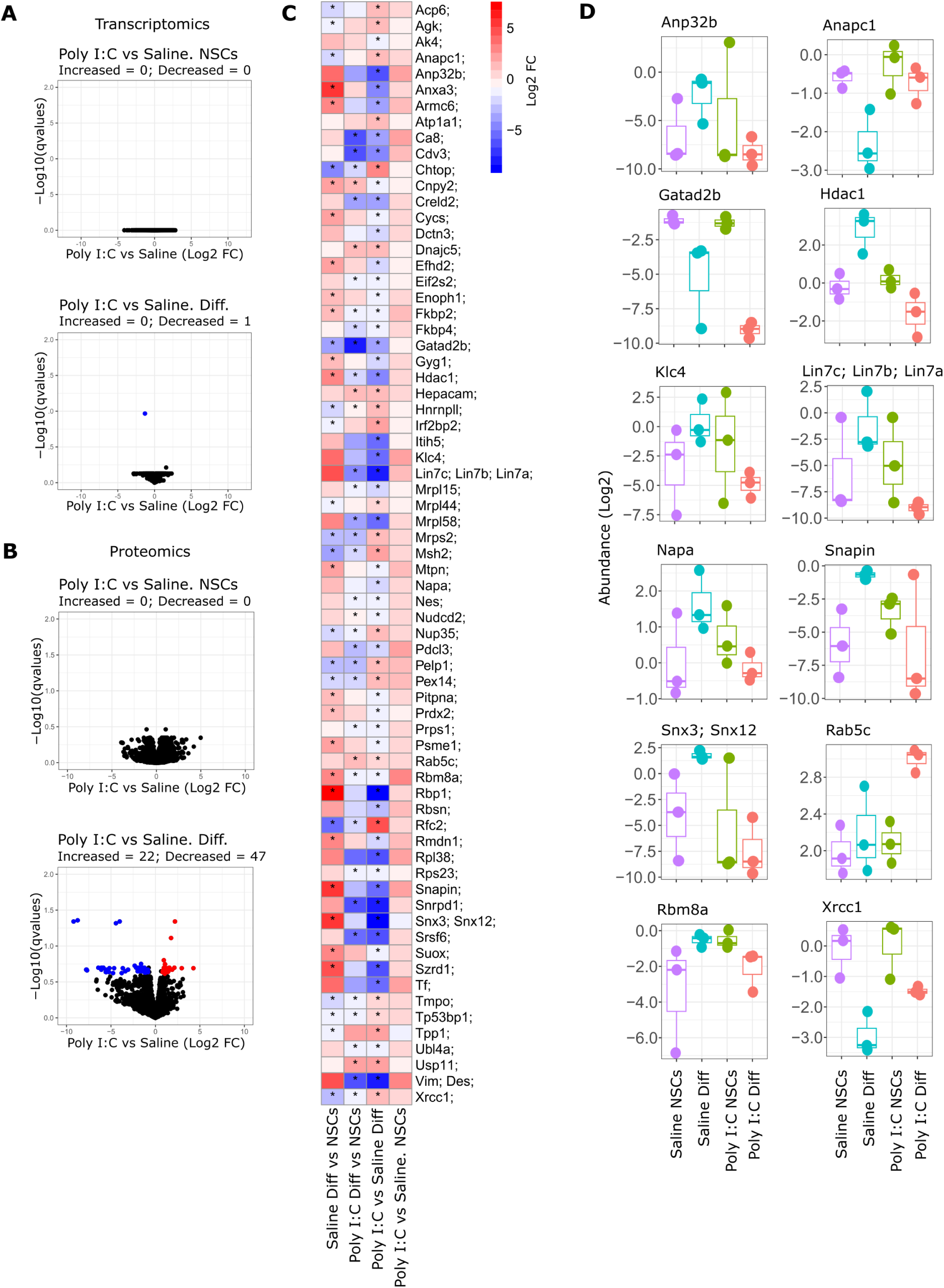
Transcriptomics and proteomics analysis comparing Poly I:C and Saline cell lines before and after differentiation of NSCs. **A, B.** Volcano plots showing transcriptomics (**A**) and proteomics (**B**) data from NSCs and differentiated cells. X-axis shows relative change expressed as Log2 Fold Change (FC) of Poly I:C versus Saline NSCs or differentiated cells. Y-axis show statistical significance expressed as -Log10 qvalues (FDR). Red and blue dots correspond to genes or proteins that changed significantly respect to Saline group (qvalues<0.25 & pval <0.05 and Log2 FC<-0.8 or Log2 FC> 0.8, respectively); black dots represent genes or proteins that did not match with the filtering criteria (FDR>0.25 & pval > 0.05, Log2 FC > −0.8 or Log2 FC <0.8). **C.** Heatmap showing the significant proteins from the bottom volcano in B. The scale colour indicated the relative change expressed as Log2 FC in each comparison indicated. The asterisks indicate the level of significance for each protein in the comparison indicated (FDR<0.25 & pval<0.05). **D.** Box plots showing the relative abundance (in Log2 scale) of some proteins represented in the heatmap from B.

### Differential phosphorylation pattern in differentiated Poly I:C derived NSCs

The results presented in previous sections showed that Poly I:C NSCs present differences in protein expression compared to Saline NSCs, only when these cells are differentiated into neurons and glial cells. Furthermore, the differential protein pattern expression occurs in proteins related to neuronal functions, protein transport or proteins involved in differentiation. To evaluate if Poly I:C induces changes in signalling pathway activation driven by phosphorylation and identify further differences between both types of NSCs, we performed phosphoproteomics approaches in differentiated cells from Poly I:C and Saline NSCs. The analysis of the basal phosphorylation levels revealed a different phosphorylation pattern between differentiated Poly I:C and Saline NSCs (Figure 4A). We found 746 phosphopeptides with increased phosphorylation and 222 phosphopeptides with decreased phosphorylation levels in differentiated Poly I:C NSCs compared to differentiated Saline NSCs (Figure 4A). Gene Ontology (GO) analysis of the phosphoproteomics data showed that significant phosphopeptides with increased phosphorylation are mostly related to cytoskeleton organization cellular component organization and biogenesis, as observed in the top significant terms with higher number of counts (Figure 4B, Supplementary Table 2A). No significant GO terms were found for proteins with decreased phosphorylation. Additionally, the GO analysis for Cellular Component terms showed that the most significant and populated terms are related with structures associated with synapse function, such as post-synaptic density or synapse, axon and dendrites (Figure 4C, Supplementary Table 2B). In this case we observed peptides with decreased and increased phosphorylation corresponding to those terms, but most populated in the case of increased phosphorylation. To expand the evaluation of the proteins identified in both GO analyses, we categorized the GO terms into groups related to axons, cytoskeleton, dendrites, neuron projections, and post-synapses (Figure 4D). We found that in most of the cases there was higher number of phosphopetides with increased phosphorylation in comparison to decreased phosphorylation, specifically in cytoskeleton, dendrite and post-synapse terms (Figure 4D). Of note, we found that proteins involved in GTPases signalling (RhoA, Ras, Cdc42 and Rac1) presented different levels of phosphorylation when comparing differentiated Poly I:C NSCs to differentiated Saline NSCs. For example, PAK2 downstream of CDC42 and RAC1, showed higher phosphorylation on S197 in differentiated Poly I:C NSCs (Figure 4E). In addition, cortactin (CTTN) also displayed altered phosphorylation on S407 and Y421 in differentiated Poly I:C NSCs (Figure 4E). Catenin Delta-1 (Ctnnd1, also known as Cadherin-associated Src substrate (CAS) or p120 catenin) also presented higher phosphorylation on S920 in differentiated Poly I:C NSCs (Figure 4E). Proto-oncogene tyrosine-protein kinase Src, also displayed higher phosphorylation on S21 in differentiated Poly I:C NSCs (Figure 4E). Drebrin (Dbn1), involved in cytoskeleton dynamics [29] also showed increased phosphorylation levels on S651 in differentiated Poly I:C NSCs (Figure 4E). Other examples of differential phosphorylation between both groups are shown in Figure 4E and in supplementary Table 3. Together, the phosphoproteomics data suggest that Poly I:C treatment induces profound changes in the activity of signalling pathways driven by protein phosphorylation.

**Figure 4.**
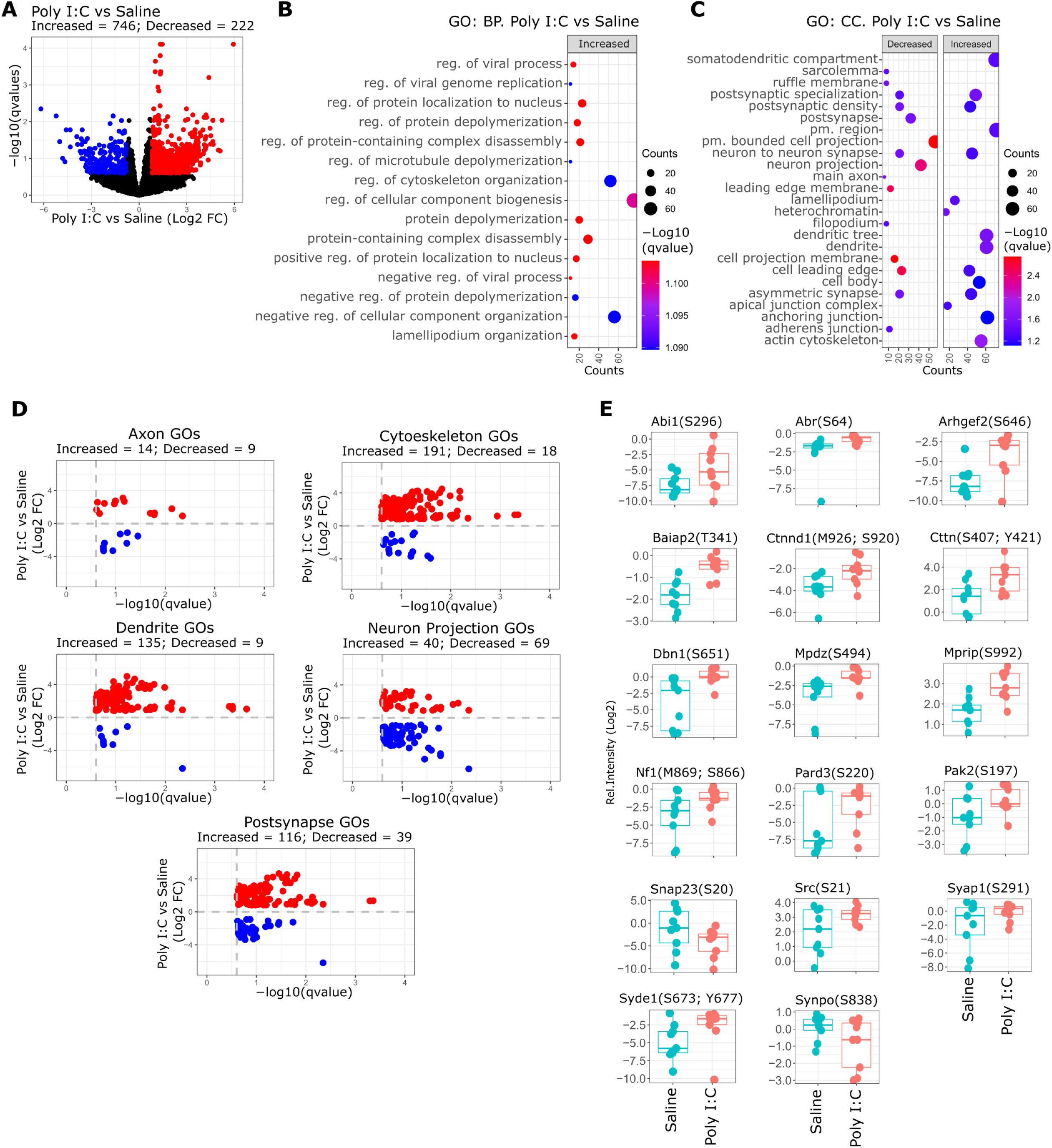
Phosphoproteomics analysis of differentiated Poly I:C and Saline NSCs. **A.** Volcano plot showing identified phosphopeptides when comparing differentiated Poly I:C versus Saline cells. X-axis shows relative change expressed as Log2 Fold Change (FC) of Poly I:C versus Saline differentiated cells. Y-axis show statistical significance expressed as -Log10 qvalues (FDR). Red and blue dots correspond to phosphopeptides that changed significantly respect to Saline group (qvalues<0.25 & pval <0.05 and Log2 FC<-0.8 or Log2 FC> 0.8, respectively), while black dots represent phosphopeptides that did not match with the filtering criteria (FDR>0.25 & pval > 0.05, Log2 FC >− 0.8 or Log2 FC <0.8). **B, C.** Gene Ontology (GO) analysis for Biological Processes (**B**) and Cellular Component (**C**) performed on the significant phosphopeptides identified in A. Plots show the top 5 most significant terms. Decreased and increased indicates GO terms which phosphopeptides were found with significant decreased or increased phosphorylation levels (blue and red dots in volcano plots). The size of the dots represents the number of proteins identified for each term, while the gradient colour scale represents the level of significance, indicated as -Log10 qvalue. **D.** Inverted volcano plots that show the phosphopeptides associated to the previous GO terms. X-axis indicates the statistical significance expressed as -Log10 of qvalues (FDR), and Y-axis represent the relative change expressed as Log2 FC of differentiated Poly I:C versus Saline cells. Red and blue dots indicate phosphopeptides with increased and decreased phosphorylation, respectively. **E.** Box plots showing some of the phospheptides identified in panel D. Y-axis show the relative intensity of the phosphopeptides in Log2 scale.

### Microtubule associated proteins (MAPs) present altered phosphorylation in differentiated Poly I:C NSCs

The analysis of proteins with altered phosphorylation pattern in differentiated Poly I:C NSCs suggest an alteration in cytoskeleton organization, which plays an essential role in neuronal function modulating the intracellular trafficking of signal transduction, as well as proper axon guidance and synapse formation [30]. To test this conjecture, we focused on the analysis of proteins involved in cytoskeleton organization and we found a different pattern of phosphorylation in the family of microtubule associated proteins (MAPs). In. particular, we observed an altered phosphorylation status in almost all members of the MAP family, with Map1a, Map1b, Map2, and Map4 exhibiting a high number of phosphopeptides (Figure 5A, top pannels). In the case of other members such as Tau (Mapt), Doublecortin (Dcx) and Doublecortin-like kinase (Dclk1), we found mostly decreased phosphorylation (Figure 5A, bottom panels and Figure 5B), but the number of altered phosphopeptides was considerably lower compared to Map2, Map4 and Map1a/b members. MAPs proteins play an important role in microtubule organization and remodelling during neuronal development, activity and maintenance, and their function are regulated by phosphorylation [31]. Our results showed an alteration in the phosphorylation of MAPs family on differentiated Poly I:C NSCs, suggesting an impairment in its function. For example, Map2, showed increased and decreased phosphorylation in 13 and 4 peptides, respectively (Figure 5A). Interestingly, we also observed alteration in the phosphorylation of the kinases Mark2 and Mark3 (Figure 5C), showing increased phosphorylation on both kinases in residues S592 (Mark2), and S543 and S543 (Mark3). Mark2/3 kinases are involved in the phosphorylation of members of MAPs family such as Map1b, Map2 and Map4, and modulate its affinity to microtubules through its phosphorylation [31].

**Figure 5.**
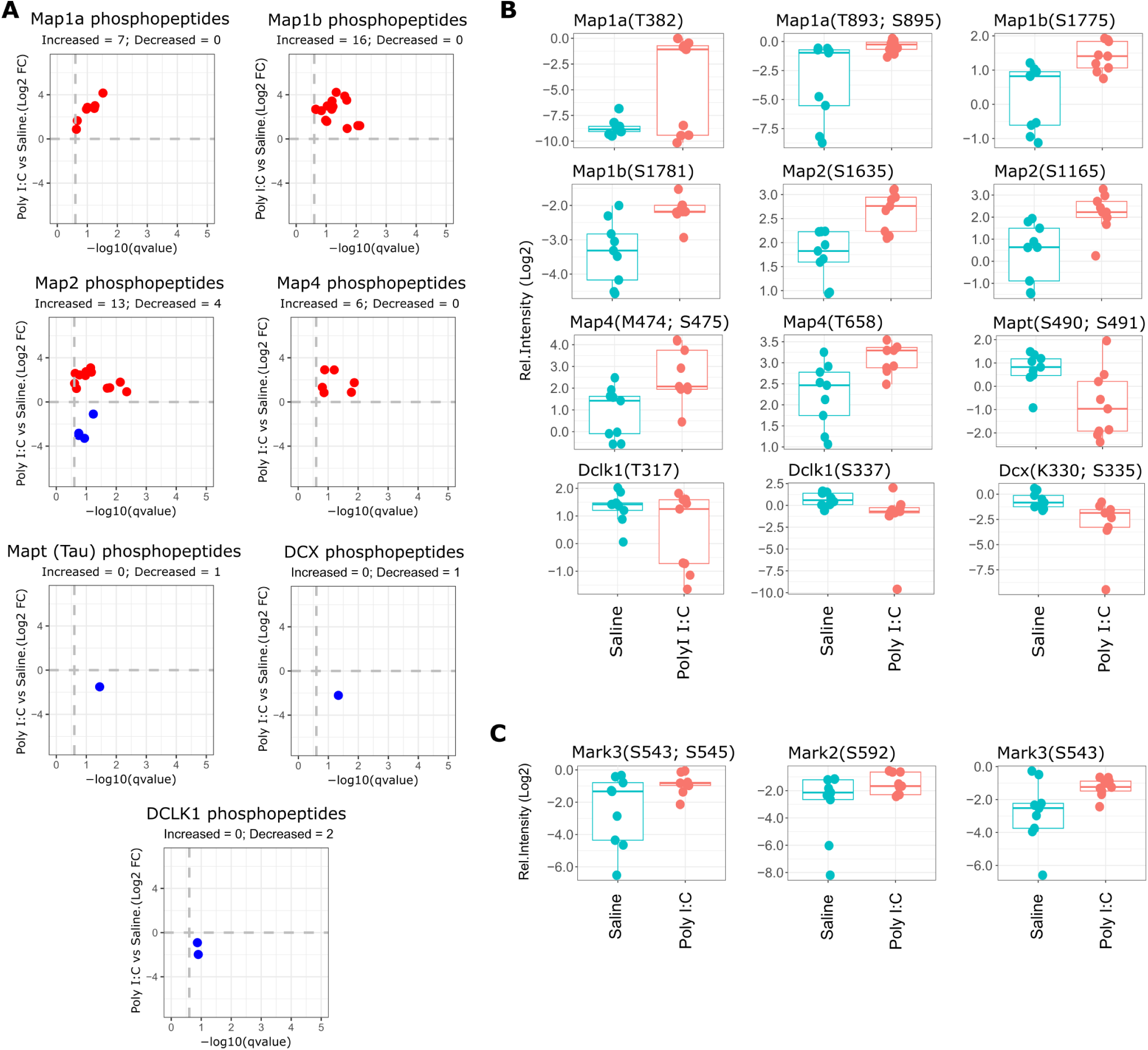
Phosphoproteomics analysis of MAPs family members. **A.** Inverted volcano plots that show the phosphopeptides identified for members of MAPs family (Map1a, Map1b, Map2, Map4, Tau, Dcx and Dclk1). X-axis indicates the statistical significance expressed as -Log10 of qvalues (FDR), and Y-axis represent the relative change expressed as Log2 FC of differentiated Poly I:C versus Saline. Red and blue dots indicate phosphopeptides with increased and decreased phosphorylation, respectively. **B.** Box plots showing some of the phosphopeptides identified in panel A. Y-axis show the relative intensity of the phosphopeptides in Log2 scale. **C.** Box plots showing the significant phosphopeptides identified for the kinases Mark3 and Mark2. Y-axis show the relative intensity of the phosphopeptides in Log2 scale.

### Functional alterations in Poly I:C NCSs

Our results showing alterations in the phosphorylation pattern of members of MAPs family, specifically Map2, imply a potential alteration in neuronal function and plasticity in differentiated Poly I:C NSCs. To test this hypothesis, we first evaluated whether the differences in phosphorylation levels in Map2 were related to different protein expression in Saline and Poly NSCs before and after its differentiation. No relevant differences were observed in protein levels in any of the groups analysed as determined by LC-MS/MS data (Figure 6A). Furthermore, no differences were observed in immunocytochemistry for Map2 in differentiated neurons from Saline and Poly I:C NSCs (Figure 6B). Interestingly, immunoblot analyses for Map2 showed a complete absence of specific immunoreactivity in differentiated Saline NSCs, accompanied by lower levels of Map2 in Poly I:C NSCs (Figure 6C). We then investigated whether differentiated neurons from Poly I:C NSCs exhibited morphological and functional alterations, as connoted by the phosphoproteomics data. To do so, we evaluated the dendritic arborization complexity in neurons from both experimental groups and found no significant differences between them (Figure 6D). However, we observed differences in dendritic spine density among differentiated neurons, with increased spine density in differentiated Poly I:C NSCs compared to differentiated Saline NSCs (Figure 6E). Additionally, we assessed the establishment of synapses among in vitro cultured neurons using a rabies virus monosynaptic tracing methodology. Interestingly, we observed a decrease in established synapses in neurons derived from Poly I:C NSCs compared to Saline neurons (Figure 6F). These data indicate that the changes in protein phosphorylation, as revealed by our phosphoproteomic data, are of biological relevance and result in an alteration in neuronal function in Poly I:C neurons.

**Figure 6.**
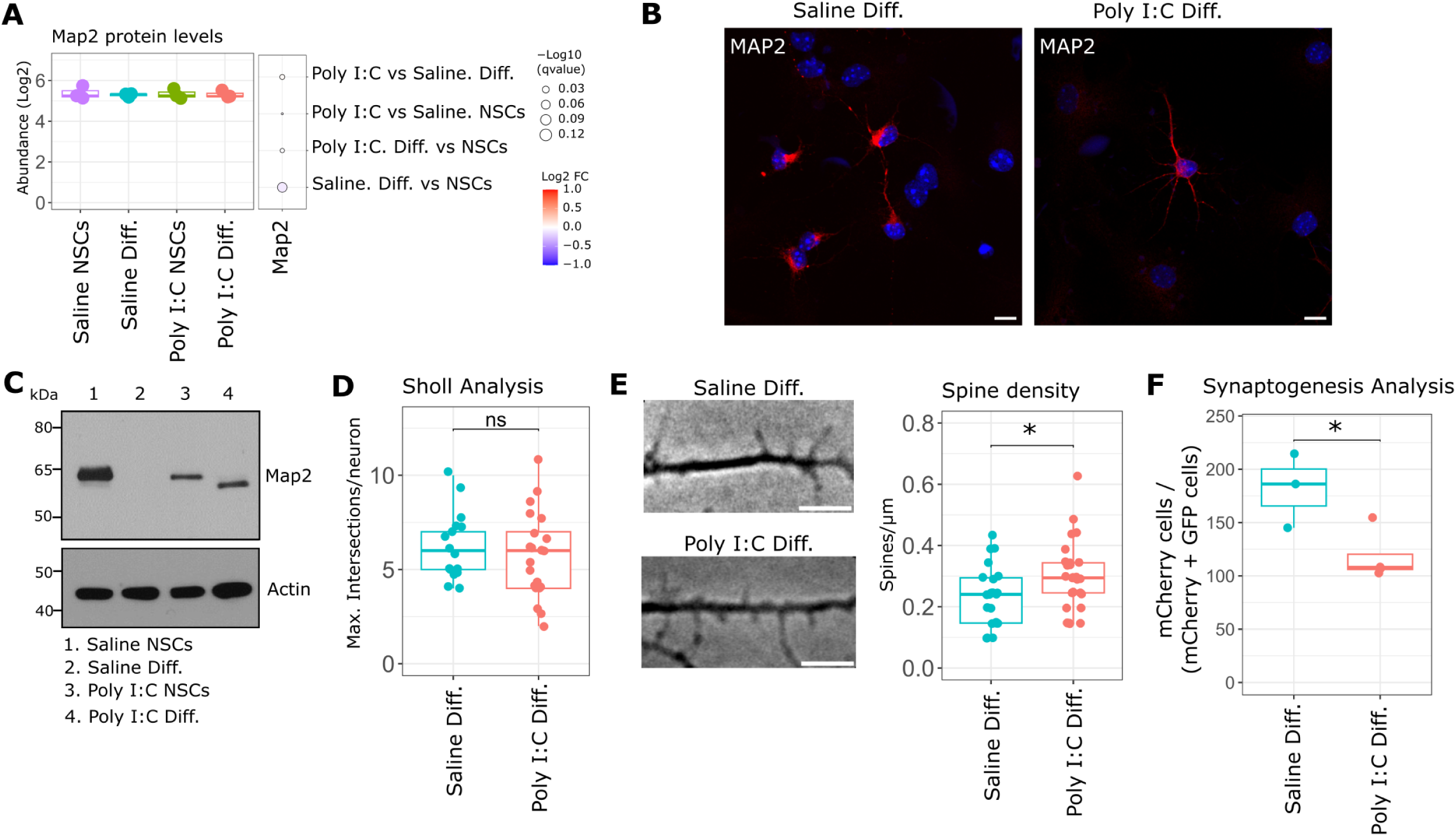
**A.** Box plots showing Map2 protein levels identified by LC-MS/MS in each of the groups analysed. Y-axis show the abundance of protein levels in Log2 scale. The dot plot at the right of the box plot indicates the relative change expressed as Log2 FC in each of the comparisons. The colour scale indicates the degree of change, increased or decreased; and the size of the dots represent the statistical significance of each comparison expressed as -Log10 qvalues (FDR). **B.** Immunocytochemistry representative images of differentiated Poly I:C and Saline NSCs for MAP2 (red) and nuclear staining with Hoechst 33342 (blue). Scale bar = 10 μm. **C.** Representative western blots for MAP2 in Saline and Poly I:C NSCs and differentiated cells. Actin was used as loading control. **D.** Box plot showing Sholl analysis for Saline and Poly I:C neurons. Each dot represents the maximum number of intersections per neuron in each condition. N = 24–30 neurons from 3 independent experiments. Error bars are SEM. **E.** Spinogenesis were evaluated by counting the number of dendritic spines in 20 µm segments of neurons from Saline and Poly I:C groups. Representative images from 20 µm segments from Saline and Poly I:C groups. Scale bar: 5 µm. Box plot shows spine densities calculated as spines/µm. N = 20–26 neurons from 3 independent experiments; error bars are SEM. **F.** Box plots showing the mean of synaptogenesis analysis in neurons from Saline and Poly I:C groups. Y-axis represent the percentage of red cells (mCherry positive) compared to the percentage of simultaneous red and green cells (mCherry and EGFP positive cells). N=3 independent experiment. Each dot represents the average of 9000-13000 cells analysed per experiment. In D, E and F, statistical differences were evaluated using unpaired T-Test. * p≤0.05, ns not significant.

## DISCUSSION

Extensive advancements and characterization of preclinical animal models have led to a deeper comprehension of the biological mechanisms of MIA in the early stages of pregnancy. These mechanisms influence the future neurodevelopment of progeny, resulting in alterations in their brain function and behaviour as they reach adulthood [1–5]. For instance, it is widely recognized at the moment that proinflammatory cytokines generated during the infection process are responsible for the effects on neurodevelopment, rather than the pathogen itself causing maternal infection [7–9]. However, there are still unknown aspects to be elucidated, such as the underlying mechanisms accountable for the emergence of neurological anomalies in offspring at a developmental stage quite distant from the immunological activation of the pregnant mother. In order to further explore into potential alterations imprinted in descendants during early gestation, we have developed an experimental cellular model based on the generation of neural stem cell lines from foetuses carried by pregnant animals exposed to MIA through treatment with Poly I:C. These neural progenitors are self-renewing and proliferative, constituting a permanent source of biological material for in vitro studies on neurons and glial cells obtained after induced differentiation. We have taken advantage of this property to conduct comprehensive omics-based research with these cells, including transcriptomics, proteomics, and phosphoproteomics studies. Overall, our results suggest that the differences between Poly I:C and Saline NSCs are primarily attributed to protein changes, particularly noticeable when cells have undergone differentiation into neurons and glia. Interestingly, these changes were not observed when comparing the transcriptomes of both experimental groups. However, they are apparent in proteomics, indicating possible divergences at the translational level. The differences found in the pattern of protein expression suggest that differentiated Poly I:C NSCs may exhibit impairment of proteins associated with neuronal functions compared to Saline NSCs. Additionally, significant differences were found when comparing the basal phosphoproteomics profile of differentiated Poly I:C and Saline NSCs. This resulted in an alteration of the phosphorylation pattern of proteins involved in processes crucial for the organization of neuronal structure and function in differentiated Poly I:C NSCs.

Numerous preceding investigations have extensively explored alterations in offspring transcriptomics caused as a result of MIA at different stages of development. The majority of them are systematically collected in a recent comprehensive review on MIA-driven transcriptomics and epigenomics alterations in specific offspring brain regions [10]. Out of a total of 118 studies included in the final review, the authors focused on 88 genes that were investigated by more than one study and after GO analysis the candidate genes were primarily grouped into three key functional areas: immune/stress response, neurotransmission/neuronal signaling and neurodevelopment. With respect epigenetic markers in MIA models, the number of studies conducted with Poly I:C in mouse were more notably reduced and indicate greater alterations in DNA methylation compared to histone modifications following MIA exposure [10, 32].

The analysis of our current transcriptomics data revealed no significant alterations in NSCs derived from the Poly I:C treatment group compared to the Saline group, either in their progenitor state or in the neural lineages obtained after differentiation. Conversely, we observed a substantial change in the pattern of gene expression within each NSC group when comparing non-differentiated cells to differentiated cells. Thus, as anticipated by the phenotypical changes observed in cells, transcriptomics and proteomics data analysis revealed a reduction in the expression of genes involved in cell division and translation processes, accompanied by a decrease in the expression of stem cell markers. On the other hand, genes related to cytoskeleton organization, actin-based processes, as well as neuronal and glial markers, were increased. An equivalent analysis of proteomics data comparing Poly I:C and Saline groups resulted in no significant differences in NSCs. In contrast, neurons and glia resulting from their differentiation presented alterations in the expression pattern of up to 69 proteins. A further detailed analysis of the corresponding transcripts encoding these proteins confirmed the lack of differences at the transcriptional level (Supplementary Fig. 5), suggesting therefore dysregulation of mRNA translation or proteostasis as previously described in MIA experiments using Poly I:C [9], which is considered a critical factor in the emergence of several neurological disorders [33, 34].

Among the proteins showing distinct abundance in differentiated Poly I:C NSCs, some are involved in the regulation of gene expression, such as HDAC1, a protein known for its role in epigenetic processes. HDACs are enzymes that catalyze the removal of acetyl groups, primarily from histones, thereby increasing their affinity for DNA and, in turn, decreasing DNA accessibility for transcription factors, resulting in a reduction of the transcriptional process. It has been reported that proinflammatory signaling induces HDAC1 ubiquitination and proteasomal degradation [35, 36]. In our experimental model, this reduction in HDAC1 does not correspond to a significant effect on DNA transcription, as reflected in the transcriptomics data, nor in neuronal differentiation from progenitors when compared to Saline NSCs [27]. Previous studies investigating HDACs in descendants of Poly I:C-treated animals yielded somehow contradictory results, showing an increase in enzymatic activity. One study reported increased global HDAC activity, particularly in females, without accompanying alterations in the expression levels of different HDAC isoforms [37]. Similarly, another study described significant hypoacetylation of histones H3 and H4 in various brain areas of juvenile offspring [38].

Phosphoproteomics data revealed that most of the proteins displaying differential phosphorylation are, directly or indirectly, related to cytoskeleton structure, which plays a key role in the organization of neuronal projections such as dendritic arborization, axons, and dendritic spines. These results suggest a potential alteration in the formation and maintenance of these projections in differentiated Poly I:C NSCs, as demonstrated by experiments on neuronal plasticity and synaptogenesis. Notably interesting is the case of the microtubule-associated proteins (MAPs) where a state of hyperphosphorylation was detected in differentiated Poly I:C NSCs for the subtypes MAP1, MAP2, and MAP4. The interactions of the MAPs with the microtubules of the cytoskeleton are affected by posttranslational modifications, such as phosphorylations, whose changes accompany the reorganization of the microtubule network during neuronal outgrowth, differentiation, and plasticity[31]. In the specific context of MAP2, recent studies have identified alterations in its phosphorylation status in brain samples obtained from schizophrenic patients [39]. A phosphoproteomic analysis of these samples revealed that among the 18 identified phosphopeptides associated with MAP2, 8 exhibited significant alterations. In our phosphoproteomic dataset, we identified 52 distinct phosphopeptides related to MAP2, with 17 of these showing elevated levels in differentiated Poly I:C NSCs compared to the corresponding Saline NSCs (Supplementary Table 4). Moreover, a lack of MAP2 immunoreactivity has also been described in brain samples from schizophrenic patients [40], which did not correspond with the results of MAP2 detection by proteomic techniques where no decrease in MAP2 expression was detected in those same samples [39, 41]. Intriguingly, we observed a total absence of immunoreactivity in immunoblotting experiments conducted on the samples corresponding to differentiated Saline NSCs. This finding was unpredictably not replicated in immunocytochemical staining, nor in the proteomic quantification of MAP2. If we consider Poly I:C NSCs as a cellular model of neurodevelopmental disease and compare our MAP2 data with those obtained from studies using postmortem brain samples from schizophrenic individuals and other animal models engineered to exhibit permanent alterations in this protein [39, 41], we observe that hyperphosphorylation of MAP2 is a consistent and distinctive feature across all cases. Conversely, the loss of immunoreactivity of MAP2, which we observed in control cells not subjected to MIA treatment, does not align with this pattern. Nonetheless, our findings may suggest a structural alteration in MAP2 during the differentiation process from NSCs to neurons/glial cells. On one hand, this differentiation process could account for the diminished immunoreactivity of the protein, particularly under denaturing conditions such as those encountered in immunoblotting electrophoresis. On the other hand, our data imply that the structural alteration differs between MAP2 derived from Poly I:C and Saline NSCs. The potential correlation between the hyperphosphorylation state of MAP2 and the structural alterations inferred from differences in immunoreactivity warrants further investigations. In any case, our model would support the hypothesis proposed by DeGiosio et al., which posits the potential pathogenic role of structural and/or post-translational alterations of MAP2 [42]. Thus, in our scenario, ’MAP2opathy’ would be responsible for changes in the post-transductional modification corresponding to phosphorylation, which would somehow affect neuronal plasticity processes such as spine formation and ultimately the establishment of synaptic connections. It is important to note that a previous MAP2 interactome study identifies an association of this protein with ribosomal complexes, suggesting a role for MAP2 in translational regulation [39]. This could relate to our data showing differences in proteomics and MAP2 alterations between Poly I:C and Saline NSCs.

Our functional studies with cultured neurons demonstrate a significant increase in dendritic spine formation in cells derived from Poly I:C NSCs, accompanied by a reduction in synaptic connections between these neurons. These results, which correlate with the hyperphosphorylation of MAP2 in this experimental group, could be interpreted following the hypothesis proposed by Sanchez et al. to explain the regulation of MAP2 phosphorylation during neuronal development [43]. Thus, during early stages of neuronal development, the elevated phosphorylation of MAP2 reduces its affinity to interact with microtubules in growing axonal cones and filopodia, providing a highly dynamic cytoskeleton necessary for axonal and dendritic growth. Once synaptic contacts have been established, MAP2 in a more dephosphorylated state would associate with microtubules, thereby stabilizing the cytoskeleton and neuronal processes at the synaptic junctions.

In conclusion, our cellular model based on NSCs derived from foetal tissue demonstrates that maternal immune activation (MIA) during early gestational stages imprints persistent alterations in the offspring stem cells. These alterations are predominantly reflected in dysregulation of protein translation and differential phosphorylation of MAP2, resulting in a deficit in neuronal maturation in terms of the establishment of synaptic connections. Further investigations will be necessary to elucidate the biological mechanisms underlying the basal hyperphosphorylation observed in MAP2.

## ACKNOWLEDGEMENTS

We thank M. Rosario Sepúlveda and Víctor Campa for their methodological guidance and technical assistance. This work was funded by the following grants and agencies: RTI2018-079344-BI00 (FEDER/Ministerio de Ciencia e Innovación-Agencia Estatal de Investigación). This work was supported by NIH R01MH084894 to J.G.-M.

## CONFLICT OF INTEREST

P.R.C. is cofounder and director of Kinomica Ltd. All other authors declare that they have no competing interests.

## SUPPLEMENTARY INFORMATION

Supplementary information is available at MP’s website.

## SUPPLEMENTARY INFORMATION (SI)

**Supplementary Figure 1.**
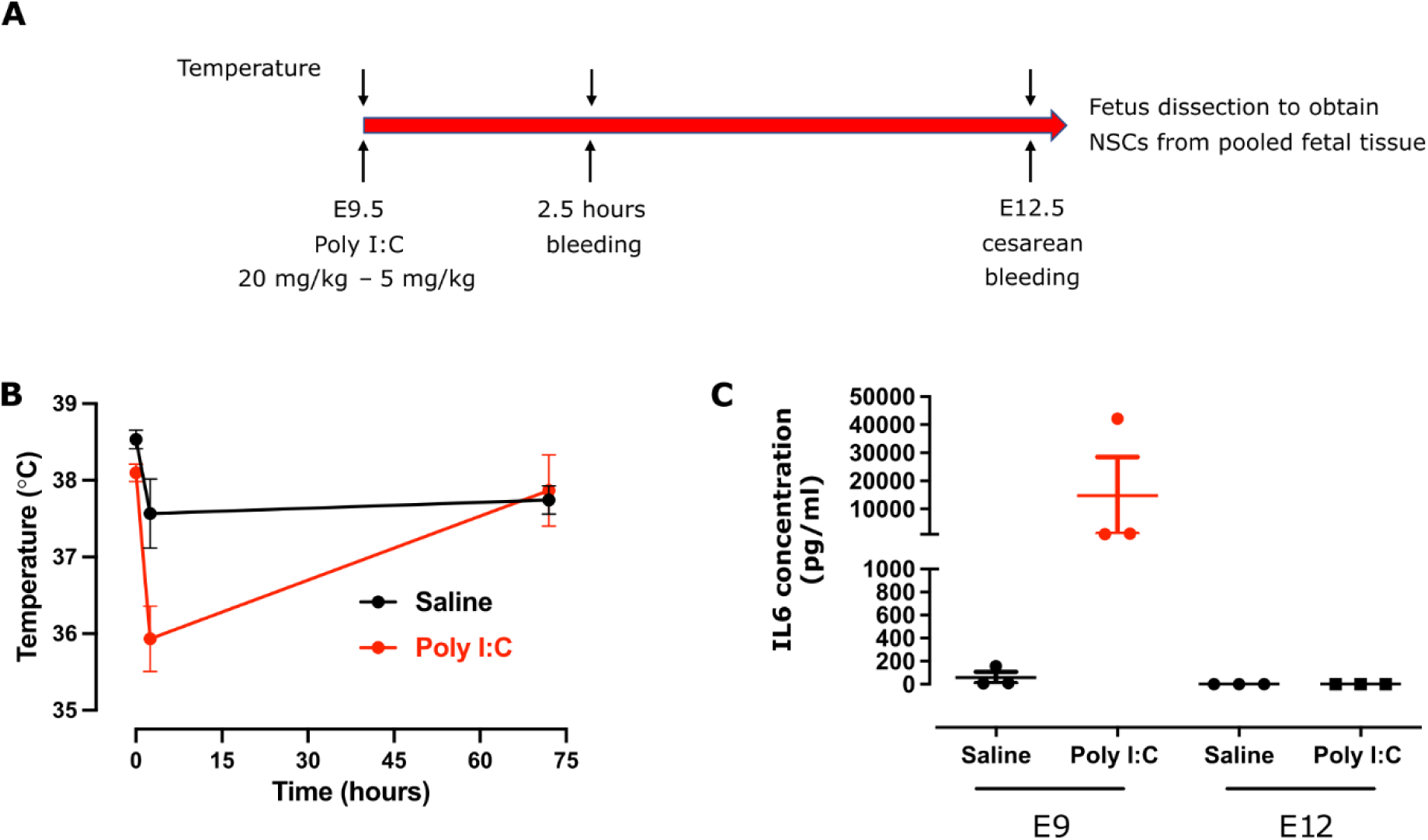
Poly I:C treatment protocol to induce maternal immune activation in pregnant mice. **A.** Timeline of the protocol from the moment of the Poly I:C acute administration to the cesarean surgery to access fetal tissue samples. Top arrowheads indicate the time points when the animal temperature was measured. Bottom arrowheads indicate the time points when blood samples were collected for subsequent IL-6 determinations. **B.** Variation in pregnant mice temperature throughout the protocol depicted in **A**. Each point represents the mean ± SEM of 3 animals. **C.** IL-6 concentration found in the serum from blood extracted from animals at the time points indicated in **A**. Each point corresponds to an individual animal, and error bars represent the SEM.

**Supplementary Figure 2.**
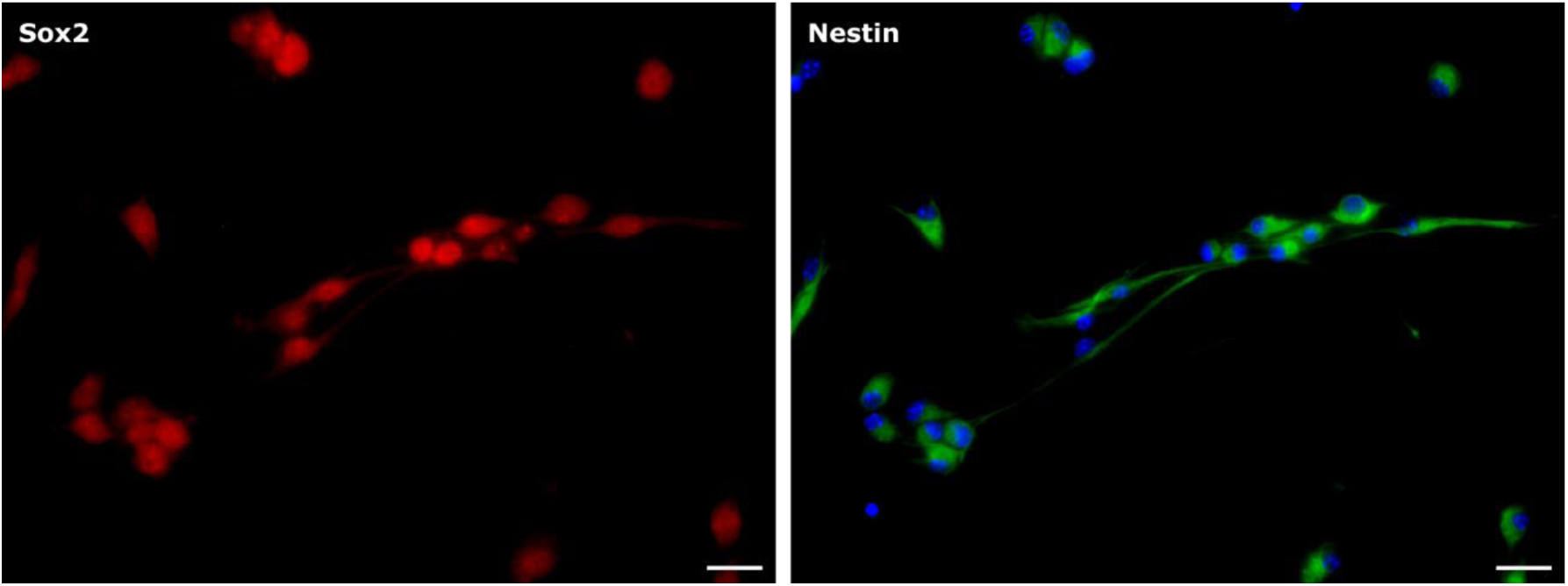
Pluripotency markers in NSCs. Representative image of NSC lines obtained from fetuses probed against pluripotency markers Sox 2(red) and Nestin (green). Cell nuclei are labelled with Hoescht 33342 (blue). Scale bar: 25 µm.

**Supplementary Figure 3.**
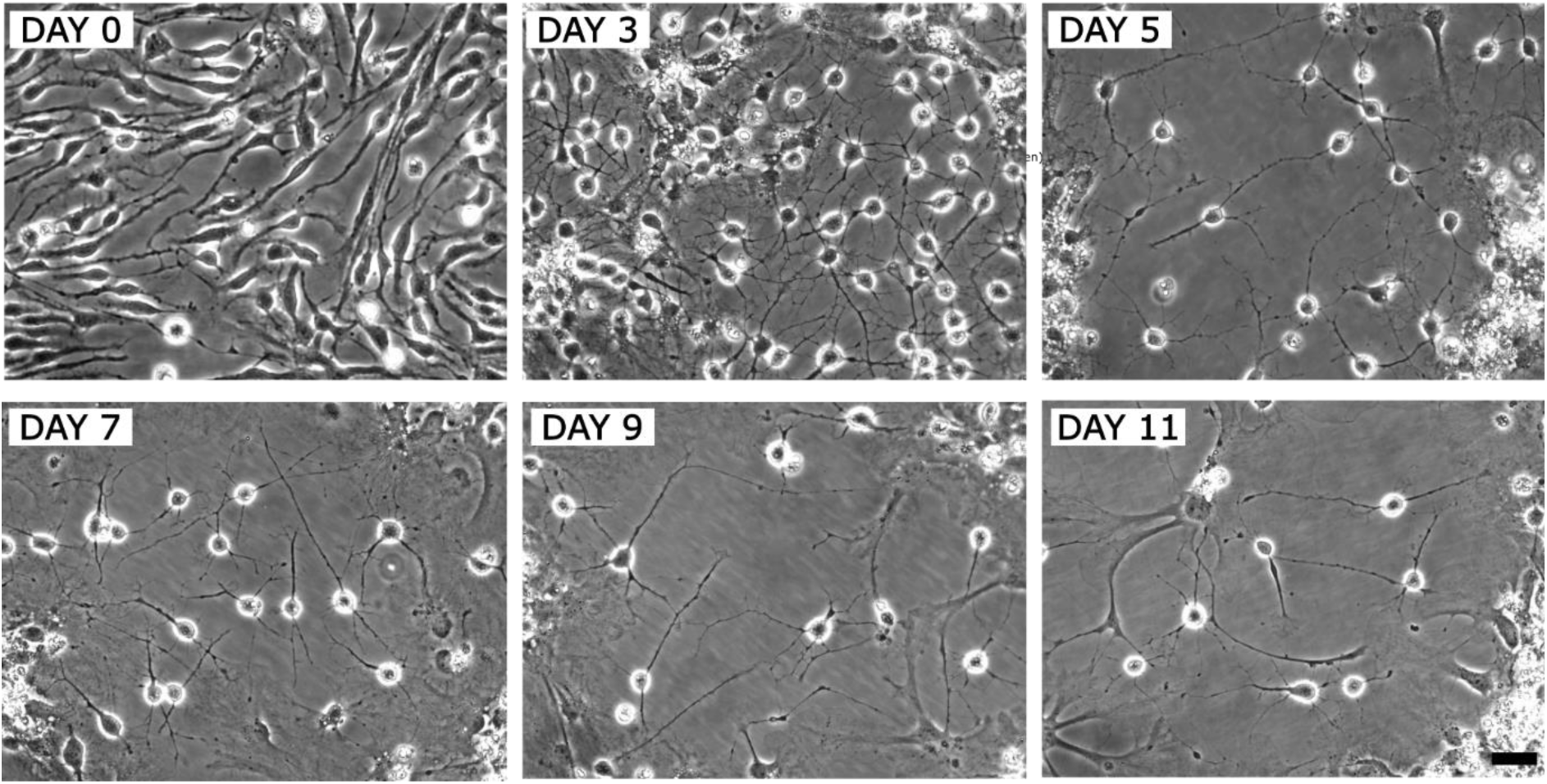
Time-lapse differentiation process of NSCs cell lines. Representative images of different time points (days of culture) during the differentiation process of NSCs. Scale bar: 50 µm.

**Supplementary Figure 4.**
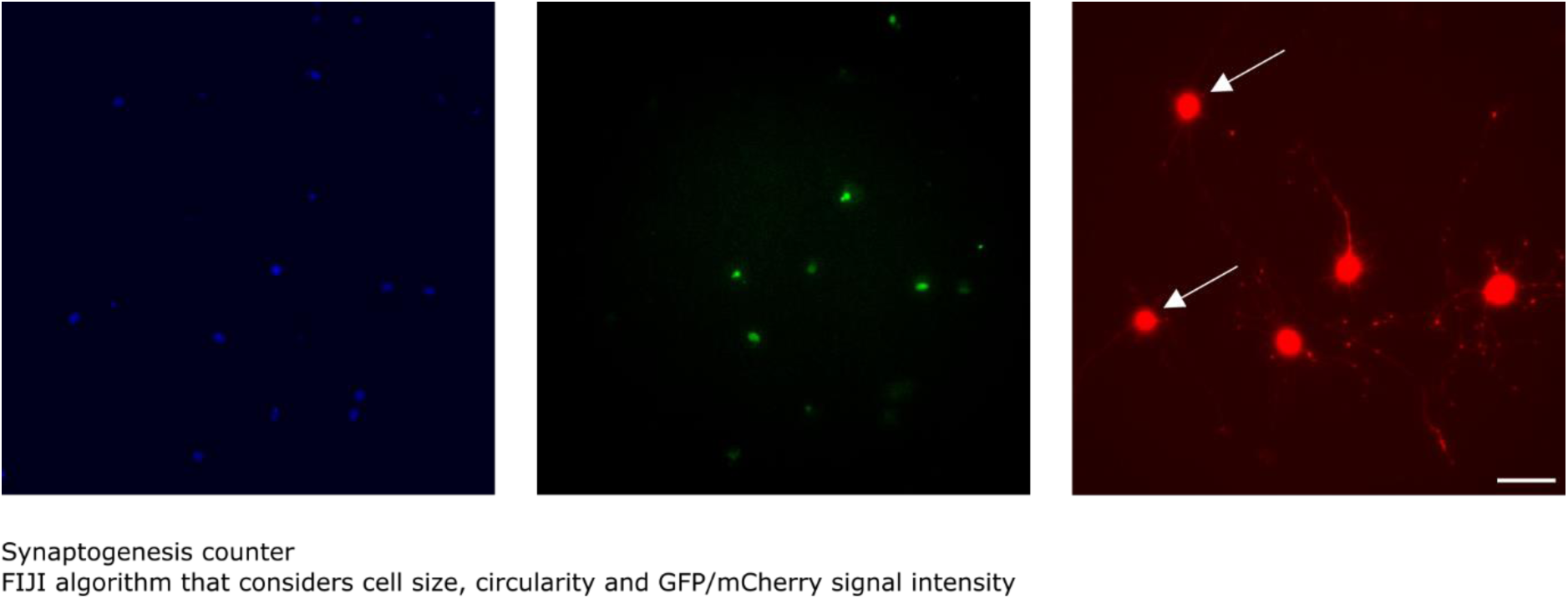
Synaptogenesis analysis using rabies virus monosynaptic tracing methodology. Representative microscopy fields are indicated by arrowheads, highlighting neurons that do not co-express green (TVA receptor) and red (recombinant rabies virus) signals. On differentiation day 3, cells were infected with lentivirus to induce the expression of the TVA receptor, along with GFP and rabies capsid glycoprotein (shown in green). Subsequently, on day 5, cells were infected with the recombinant rabies virus expressing mCherry protein (shown in red). Finally, on day 7, image acquisition from living cells was conducted after staining the cell nuclei with Hoechst 33342 (shown in blue). The nuclei staining visualizes the total population of cells within the microscopy field. Among the cells expressing the mCherry protein (red), there were cells displaying both GFP and mCherry fluorescent signals, as well as cells emitting only the mCherry signal. The image corresponding to the red channel, on the right panel, is intentionally overexposed in this case to show the dendritic arborization and thus the neuronal nature of these cells. Scale bar: 50 µm.

**Supplementary Figure 5.**
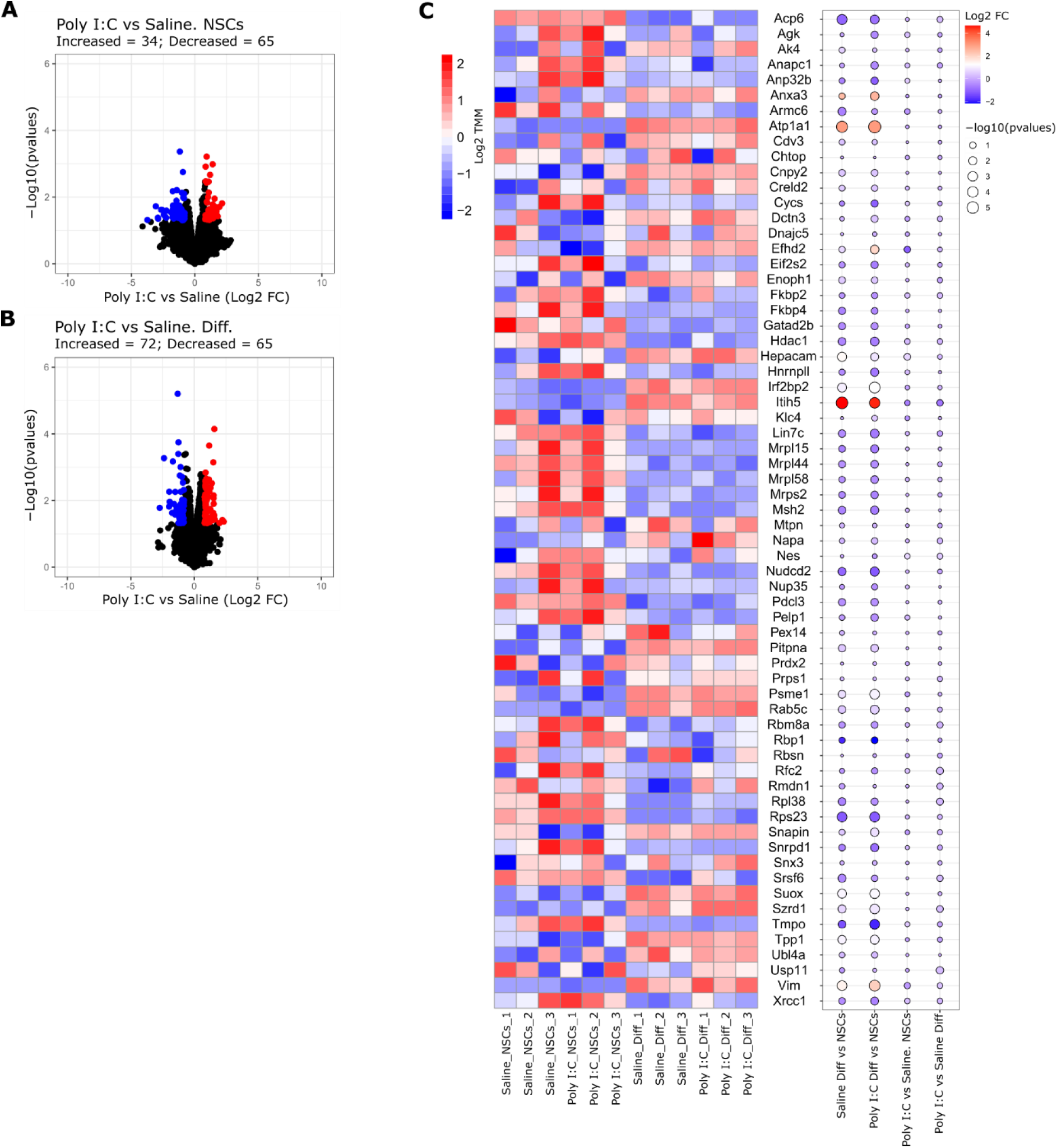
Differences in transcriptomics data between Saline and Poly I:C groups using non-adjusted p-value. Differences comparing Poly I:C and Saline before and after differentiation of NSCs. **A, B.** Volcano plots showing transcriptomics data from NSCs and differentiated cells. X-axis shows relative change expressed as Log2 Fold Change (FC) of Poly I:C versus Saline NSCs or differentiated cells. Y-axis show statistical significance expressed as -Log10 pval (non-adjusted p-value). Red and blue dots correspond to genes that changed significantly respect to Saline group (pval <0.05 and Log2 FC<-0.8 or Log2 FC> 0.8, respectively); black dots represent genes or proteins that did not match with the filtering criteria (pval > 0.05, Log2 FC > −0.8 or Log2 FC <0.8). **C**. Heatmap showing the normalized counts (expressed as Log2 TMM) in Saline and Poly I:C NSCs before and after the differentitation progress. The genes indicated in the heatmap belong to the significant proteins identified in Fig3B (bottom volcano plot) and Fig3C. Dot plot indicates the Log2 Fold change in the comparison indicated and the statistical significance (pval, non-adjusted p-value). The size of the dots indicated the level of significance for each comparison, indicated as -Log10 pvalues (pval). The colour of the dots indicates the relative change expressed as Log2 FC each comparison.

## SUPPLEMENTARY TABLES

**Supplementary table 1A. Significant Gene Ontology terms for transcriptomics data after the differentiation of Saline NSCs.**

**Supplementary table 1B. Significant Gene Ontology terms for transcriptomics data after the differentiation of Poly I:C NSCs.**

**Supplementary table 2A. Significant Gene Ontology terms for proteomics data after the differentiation of Saline NSCs.**

**Supplementary table 2B. Significant Gene Ontology terms for proteomics data after the differentiation of Poly I:C NSCs.**

**Supplementary table 3. Significant phosphopeptides identified after Gene Ontology Analysis in Fig. 4B when comparing Poly I:C versus Saline cells after the differentiation process.**

**Supplementary table 4. Identified Map2 phosphopetides when comparing Poly I:C versus Saline cells after the differentiation process.**

## REFERENCES

1. Estes ML, McAllister AK. Maternal immune activation: Implications for neuropsychiatric disorders. Science 2016; 353(6301): 772–777.

2. Meyer U. Prenatal poly(i:C) exposure and other developmental immune activation models in rodent systems. Biol Psychiatry 2014; 75(4): 307–315.

3. Meyer U, Feldon J. To poly(I:C) or not to poly(I:C): advancing preclinical schizophrenia research through the use of prenatal immune activation models. Neuropharmacology 2012; 62(3): 1308–1321.

4. Haddad FL, Patel SV, Schmid S. Maternal Immune Activation by Poly I:C as a preclinical Model for Neurodevelopmental Disorders: A focus on Autism and Schizophrenia. Neurosci Biobehav Rev 2020; 113: 546–567.

5. Reisinger S, Khan D, Kong E, Berger A, Pollak A, Pollak DD. The poly(I:C)-induced maternal immune activation model in preclinical neuropsychiatric drug discovery. Pharmacol Ther 2015; 149: 213–226.

6. Meyer U, Nyffeler M, Engler A, Urwyler A, Schedlowski M, Knuesel I et al. The time of prenatal immune challenge determines the specificity of inflammation-mediated brain and behavioral pathology. J Neurosci 2006; 26(18): 4752–4762.

7. Smith SE, Li J, Garbett K, Mirnics K, Patterson PH. Maternal immune activation alters fetal brain development through interleukin-6. J Neurosci 2007; 27(40): 10695–10702.

8. Choi GB, Yim YS, Wong H, Kim S, Kim H, Kim SV et al. The maternal interleukin-17a pathway in mice promotes autism-like phenotypes in offspring. Science 2016; 351(6276): 933–939.

9. Kalish BT, Kim E, Finander B, Duffy EE, Kim H, Gilman CK et al. Maternal immune activation in mice disrupts proteostasis in the fetal brain. Nat Neurosci 2021; 24(2): 204–213.

10. Woods RM, Lorusso JM, Potter HG, Neill JC, Glazier JD, Hager R. Maternal immune activation in rodent models: A systematic review of neurodevelopmental changes in gene expression and epigenetic modulation in the offspring brain. Neurosci Biobehav Rev 2021; 129: 389–421.

11. Kentner AC, Bilbo SD, Brown AS, Hsiao EY, McAllister AK, Meyer U et al. Maternal immune activation: reporting guidelines to improve the rigor, reproducibility, and transparency of the model. Neuropsychopharmacology 2019; 44(2): 245–258.

12. Estes ML, Farrelly K, Cameron S, Aboubechara JP, Haapanen L, Schauer JD, et al. Enhancing rigor and reproducibility in maternal immune activation models: practical considerations and predicting resilience and susceptibility using baseline immune responsiveness before pregnancy. bioRxiv 2019: 699983.

13. Pollard SM. In vitro expansion of fetal neural progenitors as adherent cell lines. Methods Mol Biol 2013; 1059: 13–24.

14. Callaway EM, Luo L. Monosynaptic Circuit Tracing with Glycoprotein-Deleted Rabies Viruses. J Neurosci 2015; 35(24): 8979–8985.

15. Osakada F, Callaway EM. Design and generation of recombinant rabies virus vectors. Nat Protoc 2013; 8(8): 1583–1601.

16. Martin-Guerrero SM, Alonso P, Iglesias A, Cimadevila M, Brea J, Loza MI et al. His452Tyr polymorphism in the human 5-HT(2A) receptor affects clozapine-induced signaling networks revealed by quantitative phosphoproteomics. Biochem Pharmacol 2021; 185: 114440.

17. Casado P, Wilkes EH, Miraki-Moud F, Hadi MM, Rio-Machin A, Rajeeve V et al. Proteomic and genomic integration identifies kinase and differentiation determinants of kinase inhibitor sensitivity in leukemia cells. Leukemia 2018; 32(8): 1818–1822.

18. Ritchie ME, Phipson B, Wu D, Hu Y, Law CW, Shi W, Smyth GK. limma powers differential expression analyses for RNA-sequencing and microarray studies. Nucleic Acids Res 2015; 43(7): e47.

19. Andres-Leon E, Nunez-Torres R, Rojas AM. miARma-Seq: a comprehensive tool for miRNA, mRNA and circRNA analysis. Sci Rep 2016; 6: 25749.

20. Andres-Leon E, Rojas AM. miARma-Seq, a comprehensive pipeline for the simultaneous study and integration of miRNA and mRNA expression data. Methods 2019; 152: 31–40.

21. Dobin A, Davis CA, Schlesinger F, Drenkow J, Zaleski C, Jha S et al. STAR: ultrafast universal RNA-seq aligner. Bioinformatics 2013; 29(1): 15–21.

22. Liao Y, Smyth GK, Shi W. featureCounts: an efficient general purpose program for assigning sequence reads to genomic features. Bioinformatics 2014; 30(7): 923–930.

23. Nikolayeva O, Robinson MD. edgeR for differential RNA-seq and ChIP-seq analysis: an application to stem cell biology. Methods Mol Biol 2014; 1150: 45–79.

24. Robinson MD, Oshlack A. A scaling normalization method for differential expression analysis of RNA-seq data. Genome Biol 2010; 11(3): R25.

25. Garbett KA, Hsiao EY, Kalman S, Patterson PH, Mirnics K. Effects of maternal immune activation on gene expression patterns in the fetal brain. Transl Psychiatry 2012; 2(4): e98.

26. Shi J, Zhao Y, Wang K, Shi X, Wang Y, Huang H et al. Cleavage of GSDMD by inflammatory caspases determines pyroptotic cell death. Nature 2015; 526(7575): 660–665.

27. Nieto-Estevez V, Changarathil G, Adeyeye AO, Coppin MO, Kassim RS, Zhu J, Hsieh J. HDAC1 Regulates Neuronal Differentiation. Front Mol Neurosci 2021; 14: 815808.

28. Cai Q, Lu L, Tian JH, Zhu YB, Qiao H, Sheng ZH. Snapin-regulated late endosomal transport is critical for efficient autophagy-lysosomal function in neurons. Neuron 2010; 68(1): 73–86.

29. Shan Y, Farmer SM, Wray S. Drebrin regulates cytoskeleton dynamics in migrating neurons through interaction with CXCR4. Proc Natl Acad Sci U S A 2021; 118(3).

30. Lasser M, Tiber J, Lowery LA. The Role of the Microtubule Cytoskeleton in Neurodevelopmental Disorders. Front Cell Neurosci 2018; 12: 165.

31. Ramkumar A, Jong BY, Ori-McKenney KM. ReMAPping the microtubule landscape: How phosphorylation dictates the activities of microtubule-associated proteins. Dev Dyn 2018; 247(1): 138–155.

32. Basil P, Li Q, Dempster EL, Mill J, Sham PC, Wong CC, McAlonan GM. Prenatal maternal immune activation causes epigenetic differences in adolescent mouse brain. Transl Psychiatry 2014; 4(9): e434.

33. Chen YC, Chang YW, Huang YS. Dysregulated Translation in Neurodevelopmental Disorders: An Overview of Autism-Risk Genes Involved in Translation. Dev Neurobiol 2019; 79(1): 60–74.

34. Jishi A, Qi X, Miranda HC. Implications of mRNA translation dysregulation for neurological disorders. Semin Cell Dev Biol 2021; 114: 11–19.

35. Gopal YN, Van Dyke MW. Depletion of histone deacetylase protein: a common consequence of inflammatory cytokine signaling? Cell Cycle 2006; 5(23): 2738–2743.

36. Villagra A, Sotomayor EM, Seto E. Histone deacetylases and the immunological network: implications in cancer and inflammation. Oncogene 2010; 29(2): 157–173.

37. Pujol Lopez Y, Kenis G, Stettinger W, Neumeier K, de Jonge S, Steinbusch HW et al. Effects of prenatal Poly I:C exposure on global histone deacetylase (HDAC) and DNA methyltransferase (DNMT) activity in the mouse brain. Mol Biol Rep 2016; 43(7): 711–717.

38. Tang B, Jia H, Kast RJ, Thomas EA. Epigenetic changes at gene promoters in response to immune activation in utero. Brain Behav Immun 2013; 30: 168–175.

39. Grubisha MJ, Sun X, MacDonald ML, Garver M, Sun Z, Paris KA et al. MAP2 is differentially phosphorylated in schizophrenia, altering its function. Mol Psychiatry 2021; 26(9): 5371–5388.

40. DeGiosio R, Kelly RM, DeDionisio AM, Newman JT, Fish KN, Sampson AR et al. MAP2 immunoreactivity deficit is conserved across the cerebral cortex within individuals with schizophrenia. NPJ Schizophr 2019; 5(1): 13.

41. Shelton MA, Newman JT, Gu H, Sampson AR, Fish KN, MacDonald ML et al. Loss of Microtubule-Associated Protein 2 Immunoreactivity Linked to Dendritic Spine Loss in Schizophrenia. Biol Psychiatry 2015; 78(6): 374–385.

42. DeGiosio RA, Grubisha MJ, MacDonald ML, McKinney BC, Camacho CJ, Sweet RA. More than a marker: potential pathogenic functions of MAP2. Front Mol Neurosci 2022; 15: 974890.

43. Sanchez C, Diaz-Nido J, Avila J. Phosphorylation of microtubule-associated protein 2 (MAP2) and its relevance for the regulation of the neuronal cytoskeleton function. Prog Neurobiol 2000; 61(2): 133–168.

